# Insulin-dependent maturation of newly generated olfactory sensory neurons after injury

**DOI:** 10.1101/2021.02.21.432129

**Authors:** Akihito Kuboki, Shu Kikuta, Nobuyoshi Otori, Hiromi Kojima, Ichiro Matsumoto, Johannes Reisert, Tatsuya Yamasoba

## Abstract

Loss of olfactory sensory neurons (OSNs) after injury to the olfactory epithelium (OE) triggers the generation of OSNs that are incorporated into olfactory circuits to restore olfactory sensory perception. This study addresses how insulin receptor-mediated signaling affects the functional recovery of OSNs after OE injury. Insulin levels were reduced in mice by ablating the pancreatic beta cells via streptozotocin injections. These streptozotocin-induced diabetic and control mice were then intraperitoneally injected with the olfactotoxic drug methimazole to selectively ablate OSNs. The OE of diabetic and control mice regenerated similarly until day 14 after injury. Thereafter, the OE of diabetic mice contained fewer mature and more apoptotic OSNs than control mice. Functionally, diabetic mice showed reduced electro-olfactogram responses and their olfactory bulbs had fewer c-Fos-active cells following odor stimulation, as well as performed worse in an odor-guided task compared to control mice. Insulin administered intranasally during day 8 to 13 after injury was sufficient to rescue recovery of OSNs in diabetic mice compared to control levels, while insulin administration between days 1 – 6 did not. During this critical time window on day 8 – 13 after injury, insulin receptors are highly expressed and intranasal application of insulin receptor antagonist inhibits regeneration. Furthermore, an insulin-enriched environment could facilitate regeneration even in non-diabetic mice. These results indicate that insulin facilitates the regeneration of OSNs after injury and suggest a critical stage during recovery (8 – 13 days after injury) during which the maturation of newly generated OSNs is highly dependent on and promoted by insulin.

**Significance Statement:** Although insulin receptor signaling is known to influence on cellular processes such as proliferation and apoptosis, it is poorly understood whether the insulin influences the regeneration of olfactory sensory neurons (OSNs) after injury. We compared the maturation processes of new OSNs after the methimazole-induced loss of pre-existing OSNs between diabetic and control mice. The results show that the regeneration of new OSNs depend on sufficient insulin levels during a specific temporal window, when insulin receptor expression is highly upregulated. Furthermore, an insulin-enriched environment via nasal insulin application during the critical period facilitates OSNs regeneration even in non-diabetic mice. The present results have implications for intranasal application of insulin as potential clinical therapeutics to facilitate OSNs regeneration after the injury.

## Introduction

Tissue homeostasis in the nervous system is a fundamental mechanism for maintaining normal function and signal transmission within complicated neural networks, such as the olfactory system. The olfactory epithelium (OE) inside the nasal cavity harbors the olfactory sensory neurons (OSNs), which are directly responsible for detecting odors. A special property of OSNs is their ability to regenerate from progenitor cells throughout life. In the OE, the ongoing incorporation of new OSNs is required to maintain the integrity of olfactory neuronal circuits for continuous monitoring of the external odor world.

Insulin is a peptide hormone produced from pancreatic beta cells and is mainly involved in regulating glucose metabolism in the periphery. Insulin and insulin receptors are also present in the central nervous system, where they regulate neuronal growth, survival, proliferation, and differentiation (Wan et al., 1997; Aberg et al., 2000; Skeberdis et al., 2001; Chiu and Cline, 2010; Fernandez and Torres-Aleman, 2012). Thus, insulin signaling in the central nervous system is of significant interest because of the global increase in the incidence of diabetes mellitus as well as the associated metabolic and neuronal comorbidities (Blazquez et al., 2014).

Insulin receptors and the corresponding mRNA are expressed in the OE (Fernandez and Torres-Aleman, 2012; Saraiva et al., 2015), where insulin profoundly affects the survival and activity of neurons. For example, insulin receptor activation increases the proliferation of cultured OSNs *in vitro* (McEntire and Pixley, 2000), enhances the electrical activity of OSNs (Savigner et al., 2009), and prevents the apoptosis of adult OSNs following bulbectomy (Lacroix et al., 2011). Thus, insulin is an endogenous factor that strongly influences the cell function and homeostasis of OSNs in the OE. However, in which cells in the OE the insulin receptor is expressed and how insulin signaling affects the proliferation and incorporation of new OSNs within olfactory neuronal circuits are poorly understood.

The OE contains primarily mature OSNs with an average lifespan of around 90 days, such that the OSN regeneration rate is low under normal physiological conditions (Benson et al., 1984; Farbman et al., 1988; Gogos et al., 2000; Magklara and Lomvardas, 2013). However, OSNs are susceptible to injury and degeneration because the OE is directly exposed to environmental agents entering the nasal cavity. In such instances, progenitor cells promptly proliferate and differentiate into OSNs, which are subsequently incorporated into olfactory circuits (Goldberg and Barres, 2000). Thus, injury resets the stage for the differentiation of OSNs, providing an opportunity to study the kinetics of OSN differentiation and circuit integration.

To study the effects of insulin signaling on OSN regeneration, we experimentally induced diabetes mellitus in mice by injecting them with streptozotocin (STZ) to destroy the pancreatic islets of Langerhans (Furman, 2015) and reduce insulin levels. In addition, the olfactotoxic drug methimazole was administered to selectively injure OSNs without damaging the progenitor cells in the OE (Sakamoto et al., 2007). These methods enabled us to examine whether OSNs generated following injury are structurally and functionally incorporated into olfactory neuronal circuits in an insulin signal-dependent manner. We found that insulin signaling via the insulin receptor facilitated the regeneration of new OSNs, which were highly susceptible to insulin deprivation-induced apoptosis 8–13 days after the injury. These results indicate that during a critical stage newly generated OSNs are dependent on insulin signaling, and suggest that insulin signaling and the insulin receptor play a key role in the homeostatic regeneration of the OE following injury.

## Materials and Methods

### Animals

C57BL/6J (stock no. 000664) and M71-IRES-tauGFP (stock no. 006676) (<u>Bozza et al., 2002</u>) strains were purchased from the Jackson Laboratory. mOR-EG-IRES-tauGFP and I7-IRES-tauGFP mice were kindly provided by Dr. Touhara (Oka et al., 2006) and Drs. Mombaerts and Bozza, respectively (Bozza et al., 2002). I7-IRES-tauGFP, M71-IRES-tauGFP, and mOR-EG-IRES-tauGFP strains are of a C57BL/6 congenic background. Male and female mice (3 weeks- to 6 months-old) were used. All experiments were performed using procedures approved by the Experimental Animal Research Committee at the University of Tokyo and by the Monell Chemical Senses Center Institutional Animal Care and Use Committee.

### Streptozotocin administration

Pancreatic beta cells of mice were ablated by intraperitoneal (i.p.) injections of streptozotocin (STZ) (120 mg/kg of body weight; Sigma-Aldrich) dissolved in PBS or 0.9% NaCl solution (saline) for three consecutive days. Seven days after the first STZ injection, fasting blood glucose levels were measured with a glucose reader (Ascensia Diabetes Care) using blood obtained from the tail veins. Mice were considered to be diabetic if fasting blood glucose levels were ≥250 mg/dl (Ishikawa et al., 2007).

### Methimazole administration

OSNs in control and diabetic mice (STZ mice) were ablated by i.p. injection of methimazole (75 mg/kg, i.p.; Sigma-Aldrich) dissolved in saline. The mice were sacrificed after 3, 7, 14, or 28 days to observe regeneration of OE and projections of OMP^+^ OSNs to OB. Projections of GFP^+^ OSNs to OB in the I7-IRES-tauGFP, M71-IRES-tauGFP, and mOR-EG-IRES-tauGFP mice were examined 45 days after methimazole injection.

### Tissue preparation

Mice were anesthetized with avertin (12.5 mg/ml) or ketamine (60 mg/kg) and tissues were fixed by intracardiac perfusion with 4 % formaldehyde (PFA) in PBS. The olfactory bulb (OB) and nose were dissected, postfixed with 4 % PFA in PBS for 2 h, soaked in 30 % sucrose in PBS, and embedded in O.C.T. compound. The noses was decalcified in 0.45 M EDTA solution (pH 8.0) before the treatment with 30 % sucrose in PBS. Coronal and/or sagittal sections of 12 µm thickness were prepared using a cryostat, mounted onto silane-coated slide glasses, and stored at -30 °C until use. Preparations including OE and OB were dissected from C57BL/6 mice, postfixed with the same fixative for 24 h, decalcified with 10 % EDTA solution (pH 7.0), and embedded in paraffin. Coronal sections (4 µm thickness) were prepared using a microtome, collected onto silane-coated slide glasses, and stored at room temperature until use. Sections were stained with hematoxylin and eosin (HE) and high-iron diamine-Alcian blue (Muto Kagaku) and immunohistochemically as described below.

### Immunohistochemistry

Paraffin-embedded sections were deparaffinized and autoclaved at 121°C for 20 min in Target Retrieval Solution (Dako Japan Inc., S1700). Frozen sections were briefly washed in PBS and incubated with 0.5 % SDS (v/v) in PBS for 15 min to retrieve antigen. Blocking was performed with 2 % bovine serum albumin (BSA) in PBS with 0.1 % Triton X-100 for 10 min for the paraffin sections and with 5 % BSA in PBS with 0.3 % Triton for the frozen sections before overnight incubation with the following primary antibodies at 4°C in a humidified chamber: anti-OMP (goat polyclonal, 1:3,000; Wako Chemicals, 544-10001-WAKO, RRID: AB_664696), anti-GAP43 (chicken polyclonal, 1:250; Thermo Fisher Scientific, PA5-95660, RRID: AB_2807462), anti-activated caspase-3 (rabbit polyclonal, 1:500; Cell Signaling Technology Inc., 9661, RRID: AB_2341188), anti-Ki67 (rabbit monoclonal, 1:300; Lab Vision, RM-9106-S1, RRID: AB_149792), and anti-c-Fos (rabbit IgG, 1:1,000; Santa Cruz Biotechnology Inc., 2250, RRID: AB_2247211). After the overnight incubation, tissues were washed with PBS and were incubated with the following secondary antibodies for 1 h at room temperature: donkey anti-goat Alexa Fluor 488 (1:100; Invitrogen, A32814, RRID: AB_2762838), donkey anti-chicken Alexa Fluor 488 (1:250; Jackson ImmunoResearch Laboratories, 703-545-155, RRID: AB_2340375), and donkey anti-rabbit Alexa Fluor 594 (1:100; Invitrogen, A21207, RRID: AB_141637). Nuclei were detected by 4’,6-diamidino-2-phenylindole (DAPI, 0.1 μg/ml). The Histofine Simple Stain MAX-PO [R] secondary antibody system (Nichirei Biosciences, 414341F, RRID: AB_2819094) was used to detect anti-Ki67. Stained and fluorescent images were acquired using a fluorescence microscope (BZ-X710, Keyence) or a Leica TCS SP2 confocal microscope (Leica Microsystems).

### Histological quantification

Histological evaluation in the OE was done for the hematoxylin and eosin (H-E) stained and immunohistochemical images of the nasal septum and the most dorsal parts of turbinate II from both the right and left nasal cavities. The thickness of the OE was measured with ImageJ software (National Institutes of Health) as the distance from the lamina propria to the surface. Cells, in H-E stained images, located between basal and apical most areas, where immature and mature OSNs reside, were counted from at least two coronal sections 500 μm apart between the caudal and rostral regions of the OE from each of 3 mice to evaluate the regeneration of OSNs as described previously (Kikuta et al., 2015). The numbers of cells with immunoreactive signals for OMP, activated caspase-3, and Ki67 were also counted from at least two coronal sections 500 μm apart (n = 3 mice). The sample size of sections from 3 mice for the histological analysis was determined based on our previous study (Kikuta et al., 2015).

Whole glomerulus and OMP^+^ areas in a given glomerulus were measured in the entire medial region from two or three coronal sections in the middle part along the anterior-posterior axis of the OB in STZ and control mice on days 14 and 28 after methimazole-induced injury. The total area of a glomerulus was delineated by the nuclei of surrounding periglomerular cells identified by DAPI staining. The OMP^+^ area in a glomerulus was determined using ImageJ software and was defined as showing an OMP signals exceeding 2 SDs of the mean background intensity in the external plexiform layer of the OB. GFP^+^ areas in a glomerulus in mOR-EG-GFP mice with and without STZ administration 45 days after methimazole injection were determined using ImageJ as described above.

### *In situ* hybridization

*In situ* hybridization was carried out using digoxigenin-labeled antisense RNA of the insulin receptor gene (*Insr*) (Accession no. NM_010568, n.t. 490-4608) as described previously (Ohmoto et al., 2008; Yamaguchi et al., 2014) and RNAscope probes (Advanced Cell Diagnostics Hayward, CA). Digoxigenin-labeled antisense RNA was synthesized and used as a probe after fragmentation to approximately 150 bases under alkaline conditions. The PFA-fixed sections were treated with proteinase K (3 μg/ml, Thermo Fisher Scientific), postfixed with 4 % PFA, acetylated with acetic anhydride, and prehybridized with salmon testis DNA. After hybridized with the riboprobe for 40 h, sections were washed in 0.2x SSC. Prehybridization, hybridization, and wash were done at 65°C. Signals of hybridized probes were detected using alkaline phosphatase-conjugated anti-digoxigenin antibodies (Roche Diagnostics, 11093274910, RRID: AB_514497) followed by 4-nitro blue tetrazolium chloride/5-bromo-4-chloro-3-indolyl phosphate as a chromogenic substrate at room temperature overnight. RNAscope assay using Mm-Insr (Advanced Cell Diagnostics, 401011) or the negative control probe to B. subtilis dihydrodipicolinate reductase (DapB) (Advanced Cell Diagnostics, 310043) was also performed in accordance with the manufacturer’s protocol. Stained images were acquired as described above.

### Odor-induced c-Fos expression in the OB

At twenty-eight days after methimazole injection, OE-ablated mice (*n* = 4–5/group) were housed individually, supplied with deodorized air through a charcoal filter, and exposed to odors 4 h after food pellets were removed. A mixture of odorants in three categories, aldehydes (propyl aldehyde, *N*-valeraldehyde, *N*-heptylaldehyde, benzaldehyde, and perilla aldehyde), lactones (γ-butyrolactone, γ-heptalactone, δ-hexalactone, δ-nonalactone, and Y-octalactone), and esters (amyl hexanoate, b-γ-hexenyl acetate, terpinyl acetate, and isoamyl acetate) was diluted 1/10 in mineral oil, and a cotton sheet soaked with 100 μl of the diluted solution was placed in a dish in the cage twice for 1 h, with a 10 min interval between placements. After the second odor exposure, mice were anesthetized with ketamine (60 mg/kg) and their OBs were dissected as described above. c-Fos expression was detected immunohistochemically. Two coronal sections (4 μm thick) from each OB at the middle part of the anterior-posterior axis were selected. The numbers of c-Fos-positive cells in the glomerular layer were counted in each of the four regions of the OB (the dorsolateral, dorsomedial, ventrolateral, and ventromedial parts) using images taken at 20× magnification with a fluorescence microscope.

### Electro-olfactogram recordings

Electro-olfactogram (EOG) recordings were performed on STZ mice on day 90 after STZ administration and control mice on day 90 after saline administration without methimazole-induced injury, and 10-week-old control and STZ mice 14 and 28 days after methimazole-induced injury to evaluate the odor-induced response of the OE. The mice were euthanized with CO2 and decapitated, and their heads were sagittally bisected at the center of the nasal septum. The septum was removed to expose the olfactory turbinates in the nasal cavity. The bisected heads were quickly transferred to a recording setup, where a stream of humidified air flowed (3 l/min) over the tissue.

Pentyl acetate (Sigma-Aldrich) was first dissolved in DMSO to make a 5 M stock solution, which was diluted in water to obtain odorant solutions ranging from 1 × 10^−7^ to 10^−1^ M in 5 ml final volumes in sealed 50-ml glass bottles. As a control (0 M odorant), a solution of DMSO equivalent to the concentration in the 10^−1^ M odorant solution was used. The headspace from each odorant solution was injected with a Picospritzer (Parker Hannifin) into the air stream flowing over the OE to stimulate OSNs. The OE was exposed to each odorant solution and the odorant-induced responses were recorded in the following order; 10^−1^ M, 10^−2^ M, 10^−3^ M, 10^−4^ M, 10^−5^ M, 10^−6^ M, 10^−7^ M, 0 M, 10^−7^ M, 10^−6^ M, 10^−5^ M, 10^−4^ M, 10^−3^ M, 10^−2^ M, and 10^−1^ M.

To record the EOGs, two electrodes were placed on the surfaces of turbinate II and IIB from either the left or right half of the head at similar positions in control and STZ mice (Barrios et al., 2014). The signals were recorded with two DP-301 amplifiers (Warner Instruments), and the 1 kHz low-pass-filtered signal was digitized at 2 kHz with a Micro1401 mkII digitizer and Signal ver. 5.01 software (Cambridge Electronic Design). The data were analyzed using Origin software ver. 8.5 (Origin Lab). The investigator was blinded to the mouse treatment for all EOG experiments until data analysis was completed. When the odorant-induced response showed more than 25% difference between the first 10^−1^ M and the last 10^−1^ M odorant exposure, the experiment was excluded from further analysis.

### Odor-guided cookie search test

Twenty-eight days after methimazole-induced injury, a cookie test was performed for 4 consecutive days in control and STZ mice (7 mice/group) as previously reported (Stephan et al., 2011; Pietra et al., 2016). Starting at the same time of each day, the mice were food deprived for 6 h with water access *ad libitum*. Each mouse was then transferred to a standard mouse cage (length, 24 cm; height, 12 cm, width, 15 cm), in which a 7–10 mm^3^ piece of cookie (Oreo; Nabisco) had been buried under fresh bedding at a 3 cm depth on trial days 1, 2, and 3; on trial day 4, the cookie was placed on the surface of the bedding. The cookie was placed in a randomly chosen corner of the cage in every trial to prevent mice from predicting the position of the cookie on the basis of spatial information. After the mouse was placed in the cage, the latency to find the cookie was recorded, defined as the time until the mouse located the position of the cookie, dug the bedding, and bit into the cookie. The maximal time for this search task was set at 10 min. If the mouse failed to retrieve the cookie, it was exposed on the bedding for the mouse to eat. Once the trial was finished, each mouse was returned to its original cage. The test was conducted once each trial day. The order of tested mice was randomly chosen each day.

### Intraperitoneal and intranasal insulin administration

For i.p. insulin injections, insulin detemir (100 units/ml; Novo Nordisk) was injected into STZ mice on day 1–13, 1–6, or 8–13 after methimazole injection (3 units/kg per administration; see Fig. 7*A*).

For intranasal administrations (Marks et al., 2009), Humulin R (100 units/ml; Lilly USA) was applied to STZ mice with the same time course as for i.p. administration. For this, 30 μl drops of insulin diluted 1/1, 1/2, or 1/3 with saline were applied to each nostril, and the insulin solution was drawn into both nasal cavities by the animal’s natural inhalation. This was repeated three times each day for a total 90 μl of insulin solution per day. To examine the effect of the nasal insulin administration on blood glucose levels, fasting blood glucose levels at 60 and 120 min after the administration of the 1/1, 1/2, or 1/3 concentration of insulin (Fig. 7*G*) were measured with a glucose reader (Ascensia Diabetes Care).

### Unilateral intranasal insulin receptor antagonist administration

To evaluate the effect of blocking the insulin receptor in the nasal cavity, the insulin receptor antagonist, S961 (0.5 µg/µl; Phoenix Pharmaceuticals) was unilaterally applied to awake mOR-EG-GFP mice on day 8-13 after the methimazole injection. A volume of 10 µl of S961 was applied twice a day, resulting in a total 10 µg of S961 application each day. In control mice, PBS was unilaterally applied with the same time course as for S961 administration. The septal OE of three coronal sections from the caudal to rostral regions was analyzed for both the sides that received S961 or PBS and the non-applied control. To examine the effect of S961 on blood glucose levels, fasting blood glucose levels were measured at 120 min after the administration (Fig. 8*E*) with a glucose reader (Ascensia Diabetes Care).

### Statistical analysis

Statistical analyses were performed with Mann-Whitney *U* tests (Figs 1*D,E,I,J*; 2*C,D,F*; 4B,E,F,*I*; 6*B,D*; 7*C–E,I*; 8*G–I*, and 9*C,F*), Student’s *t* tests (Figs 1*K,L*; 3*A–D*; 5*C*), the Steel-Dwass test (Fig. 6*D*), and the Steel test (Fig. 7*G*). The data are presented as the means ± SDs except for the results of EOG and odor-guided cookie search test (Figs 1*L*; 3*B,D*; 5*C*, mean ± SEM). A *p* value of <0.05 was considered to be statistically significant.

## Results

### Expression of the *Insr* gene in the OE

First, we examined in which cell type the *Insr* gene is expressed in the OE by performing *in situ* hybridization on an OE of a control mice using digoxigenin-labeled antisense RNA of *Insr* (Fig. 1*A*) and RNAscope assay against *Insr* (Fig. 1*B*) (also see Fig. 8*A* for an RNAscope assay against *Insr*). Fig. 1*A* shows a representative image of a coronal OE section at low (left panel) and high magnification (middle and right two panels). Fig. 1*A,B* (left panels) shows that the mRNA signal for *Insr* is visible in the turbinates, as well as in the nasal septum. Fig. 1*A* (middle panel) shows that the *in situ* signal is observed in the apical and basolateral layers of the OE (see also Fig. 8*A*), where supporting cells and immature OSNs are located. Also note the somewhat non-homogeneous staining, with some areas showing higher levels of *Insr* expression compared with others (right panels in Fig. 1*A,B*).

**Figure 1.**
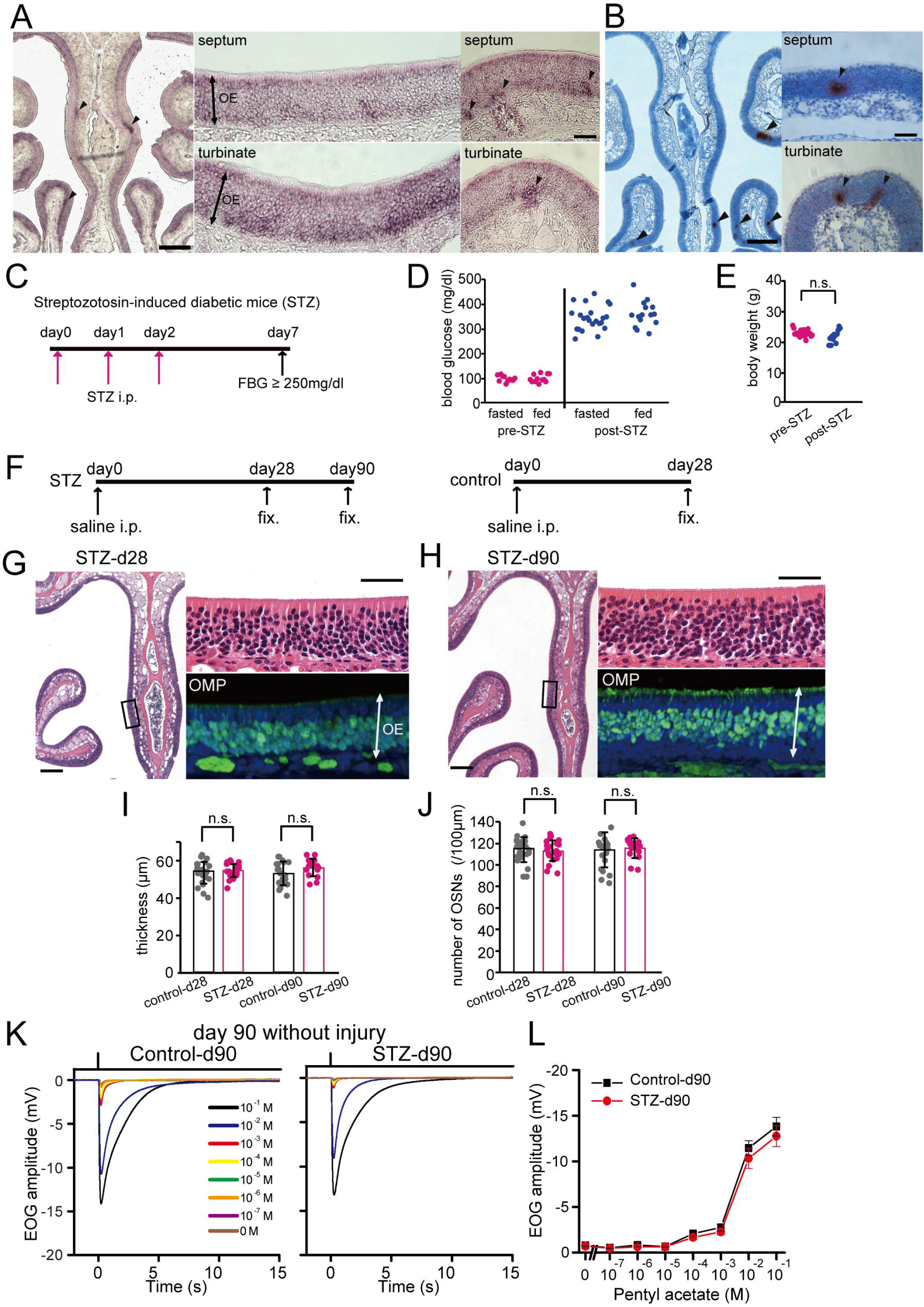
Decreased insulin for 90 days does not induce histological changes in OE. ***A, B***, *Insr* mRNA of uninjured OEs detected by *in situ* hybridization using digoxigenin-labeled antisense RNA and RNAscope probes. Representative image for an *in situ* hybridization using digoxigenin-labeled antisense RNA of *Insr* of an uninjured OE in a 3-wk-old mouse (*n* = 2 mice) (***A***) and image for an RNAscope assay against *Insr* shown as small brown dots on an uninjured OE of a 10-wk-old mouse (*n* = 2 mice) (***B***). Signals were detected especially in the apical and bottom layers of the OE, where supporting cells and immature OSNs, respectively, are located (middle panel in ***A***). Arrowheads indicate the non-homogeneous staining with higher level of *Insr* expression (***A, B***). Scale bars: 250 µm at low magnification, 50 µm at higher magnification. ***C***, Protocol of STZ treatment. STZ was intraperitoneally (i.p.) injected on days 0, 1, and 2. Mice with a fasting blood glucose (FBS) ≥250 mg/dl were considered diabetic. ***D***, Fasted and fed blood glucose levels before and after STZ administration. Both fasted and fed blood glucose levels after STZ administration (post-STZ) were much higher than those before STZ administration (pre-STZ). ***E***, Body weights before and after STZ administration. There was no significant difference (n.s.) in body weights between pre-STZ and post-STZ (Mann-Whitney *U* test). ***F***, Experiment timelines for STZ mice obtained at 28 and 90 days, and saline-injected (i.p.) control mice for comparison at 28 days. ***G, H***, Representative coronal sections of nasal septa showing the OEs stained with hematoxylin and anti-OMP antibody (green) from STZ mice on day 28 (***G***) or day 90 (***H***) after saline injection. Left images, lower magnification; right upper (hematoxylin) and lower (OMP) images, higher magnifications of the OE depicted in the left photographs indicated by squares. Scale bars: 100 µm at lower magnification, 50 µm at higher magnification. ***I, J***, Thicknesses of the OEs (***I***) and density of OSNs (***J***) in STZ-d28, STZ-d90, and saline-d28 mice. There were no significant differences between STZ-d28 and STZ-d90 or STZ-d90 and saline-d28 mice (Mann-Whitney *U* tests). ***K,*** Odorant-evoked EOG responses to pentyl acetate at different concentrations in control and STZ mice on day 90 without injury. Similar response kinetics of the EOG were observed in control and STZ mice. ***L,*** Comparison of peak amplitudes in EOG recordings between control and STZ mice (*n* = 6 mice/group; Student’s t test) on day 90 without injury. Relative to control mice, the EOG amplitudes in response to each of all concentrations of pentyl acetate were not significantly different in STZ mice (10^−1^ M odorant solution, *p* = 0.504; 10^−2^ M, *p* = 0.419; 10^−3^ M, *p* = 0.355; 10^−4^ M, *p* = 0.245; 10^−5^ M, *p* = 0.880; 10^−6^ M, *p* = 0.330; 10^−7^ M, *p* = 0.604; Student’s *t* tests). Error bars, SEM.

### Decreased insulin signaling for 90 days does not alter OE structure

To examine the effect of insulin signaling on tissue homeostasis in uninjured OE, we generated diabetic mice using i.p. injections of STZ (Fig. 1*C*). On average (± SD), both fasted and fed blood glucose levels in mice at 10 days after STZ treatment were higher [fasted, 343 ± 48 mg/dl (23 mice); fed, 355 ± 51 mg/dl (15 mice)] than before STZ administration [fasted, 94 ± 12 mg/dl (11 mice); fed, 100 ± 16 mg/dl (12 mice)] (Fig. 1*D*). We did not detect differences in the body weights of the mice before and after STZ administration [22.5 ± 1.1 g (20 mice) vs. 21.5 ± 2.1 g (12 mice), respectively, *p* = 0.080, Mann-Whitney *U* test] (Fig. 1*E*).

We next examined histological changes in the OEs of STZ mice [fixation on day 28 (STZ-d28) and 90 (STZ-d90) after STZ administration] and control mice [fixation on day 28 (saline-d28) after saline administration] (Fig. 1*F*). Fig. 1*G,H* shows representative coronal sections of OEs stained with hematoxylin (left and upper right) and immunostained with an anti-OMP antibody to identify mature OSNs (lower right) in STZ-d28 and STZ-d90 mice.

The OE thickness and the number of OSNs in STZ-d90 mice did not differ from those in STZ-d28 and saline-d28 mice (saline-d28 (*n* = 3 mice), STZ-d28 (*n* = 3 mice), saline-d90 (*n* = 3), STZ-d90 (*n* = 2 mice); OE thickness: saline-d28 vs. STZ-d28, *p* = 0.488; saline-d90 vs. STZ-d90, *p* = 0.153; number of OSNs: saline-d28 vs. STZ-d28, *p* = 0.695; saline-d90 vs. STZ-d90, *p* = 0.781; Mann-Whitney *U* tests) (Fig. 1*I,J*). To examine whether odorant-induced responses in the OE were changed in STZ-d90 and saline-d90 mice, EOG recordings were performed (*n* = 6 mice/group) (Fig. 1*K,L*). Consistent with the histological results, EOG response amplitudes to the odorant pentyl acetate were not significantly different between both groups of mice (pentyl acetate: 10^−1^ M odorant solution, *p* = 0.504; 10^−2^ M, *p* = 0.419; 10^−3^ M, *p* = 0.355; 10^−4^ M, *p* = 0.245; 10^−5^ M, *p* = 0.880; 10^−6^ M, *p* = 0.330; 10^−7^ M, *p* = 0.604; Student’s *t* tests) (Fig. 1*L*). These results suggest that decreased insulin signaling alone for 90 days does not induce histological changes and reduces odorant-induced responses in uninjured OE.

### Insulin signaling is required for the replacement of functional OSNs after OE injury

We next investigated whether decreased insulin signaling affects the incorporation of new neurons following OE injury. Methimazole, an olfactotoxic drug, activates an apoptotic cascade in OSNs throughout the OE (Brittebo, 1995; Sakamoto et al., 2007). The lost OSNs are replaced by new OSNs from proliferating progenitor cells, such that the OE returns to its preinjury state after one month (Schwob, 2002; Kikuta et al., 2015). To examine whether insulin signaling contributes to this recovery, we assessed histological changes in the OE 3, 7, 14, and 28 days after methimazole-induced injury in control and STZ mice (Fig. 2*A*). The OEs in control and STZ mice recovered similarly during the first 7 days after methimazole-induced injury, and no significant differences in the thicknesses of the OEs or the numbers of OSNs were observed (OE thickness, *p* = 0.579; numbers of OSNs, *p* = 0.160; Mann-Whitney *U* tests) (Fig. 2*B–D*). The OE thickness and number of OSNs in control mice gradually increased to match the levels observed with saline administration on day 28 (control-d28 (*n* = 3 mice) vs. saline-d28 (*n* = 3 mice): OE thickness, *p* = 0.702; numbers of OSNs, *p* = 0.796; Mann-Whitney *U* tests) (Fig. 2*C,D*). By comparison, the recovery in STZ mice was significantly suppressed [control (*n* = 3 mice) vs. STZ (*n* = 3 mice): OE thickness: day 14, *p* = 0.001, day 28, *p* = 0.002; numbers of OSNs: day 14, *p* < 0.001, day 28, *p* = 0.002; Mann-Whitney *U* tests] (Fig. 2*B–D*).

**Figure 2.**
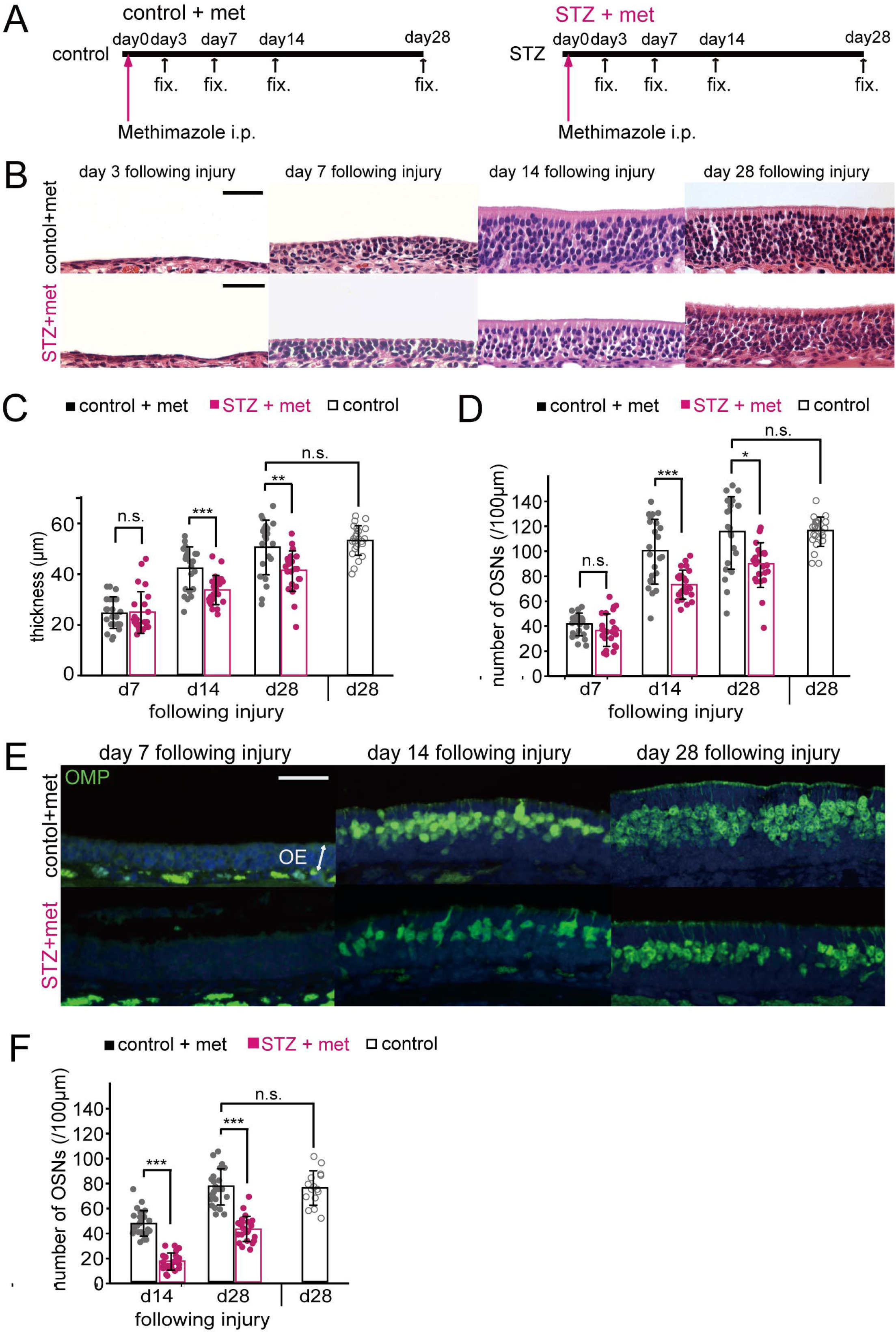
Decreased insulin disrupts OE recovery following methimazole-induced injury. ***A***, Time course of the experimental design in both control and STZ mice. Methimazole was injected (i.p.), and the tissues were fixed (“fix.”) after 3, 7, 14, and 28 days. ***B***, Representative coronal sections of the nasal septa stained with hematoxylin 3, 7, 14, and 28 days after methimazole-induced injury in both control and STZ mice. Scale bars, 50 μm. ***C, D***, The OE thicknesses (**C**) and the density of OSNs (**D**) for control, STZ, and saline-administered mice on days 7 (d7), 14 (d14), and 28 (d28) after methimazole-induced injury. On days 14 and 28, the OE thickness and density of OSNs in STZ mice were reduced significantly compared to those in control mice (* *p* < 0.05; ** *p* < 0.01; *** *p* < 0.001; Mann-Whitney *U* test). On day 28 following methimazole-induced injury, the thickness of the OEs and the density of OSNs in the control mice were restored to levels similar to those in saline-administered mice (Mann-Whitney *U* test). ***E***, Representative coronal sections stained with anti-OMP antibody (green) 7, 14, and 28 days after methimazole-induced injury in control and STZ mice. Scale bar, 50 μm. ***F***, Density of OMP-positive cells in control and STZ mice. On days 14 and 28 following methimazole-induced injury, the density of OMP-positive cells was significantly lower in STZ mice than in control mice (*** *p* < 0.001; Mann-Whitney *U* test). The density of OMP-positive cells in control mice 28 days after injury was restored to levels observed in saline-administered mice (Mann-Whitney *U* test).

To examine whether the impaired regeneration in STZ mice is also accompanied by reduced numbers of mature OSNs, we quantified the number of OMP-immunostained OSNs in the OE. Fig. 2*E* shows representative pictures of nasal septum sections stained with anti-OMP antibodies. OMP-positive cells were not detectable 7 days after injury, but emerged after 14 days in both control and STZ mice (Fig. 2*E*); however, there were significantly fewer OMP-positive cells in STZ mice than in control mice (control-d14 (*n* = 3 mice) vs. STZ-d14 (*n* = 3 mice): *p* < 0.001; Mann-Whitney *U* test) (Fig. 2*F*). On day 28, the number of OMP-positive cells in control mice matched the number in saline-treated (uninjured) mice (control-d28 (*n* = 3 mice), vs. saline-d28 (*n* = 2 mice), *p* = 0.966; Mann-Whitney *U* test) (Fig. 2*F*), whereas the number in STZ mice was significantly lower than in control mice (control-d28 vs. STZ-d28, *p* < 0.001; Mann-Whitney *U* test) (Fig. 2*F*). These results indicate that decreased insulin signaling results in incomplete recovery of the OE with fewer newly generated OSNs at 14 and 28 days after injury.

We performed EOG recordings in control and STZ mice to examine whether the incomplete recovery of the OE after STZ administration on days 14 and 28 after injury was accompanied by decreased odor-induced responses (*n* = 6 mice/group) (Fig. 3). We observed similar response kinetics in the EOG recordings from control and STZ mice (Fig. 3A, *C*). However, the amplitudes in response to higher odorant concentrations in STZ mice were significantly lower than those in control mice both at day 14 (pentyl acetate: 10^−1^ M odorant solution, *p* = 0.002; 10^−2^ M, *p* = 0.002; 10^−3^ M, *p* = 0.018; Student’s *t* tests) (Fig. 3*B*) and at day 28 (pentyl acetate: 10^−1^ M odorant solution, *p* = 0.012; 10^−2^ M, *p* = 0.015; 10^−3^ M, *p* = 0.044; Student’s *t* tests) (Fig. 3*D*). These results suggest that an incomplete recovery of the OE in STZ mice is reflected in decreased odor-evoked responses.

**Figure 3.**
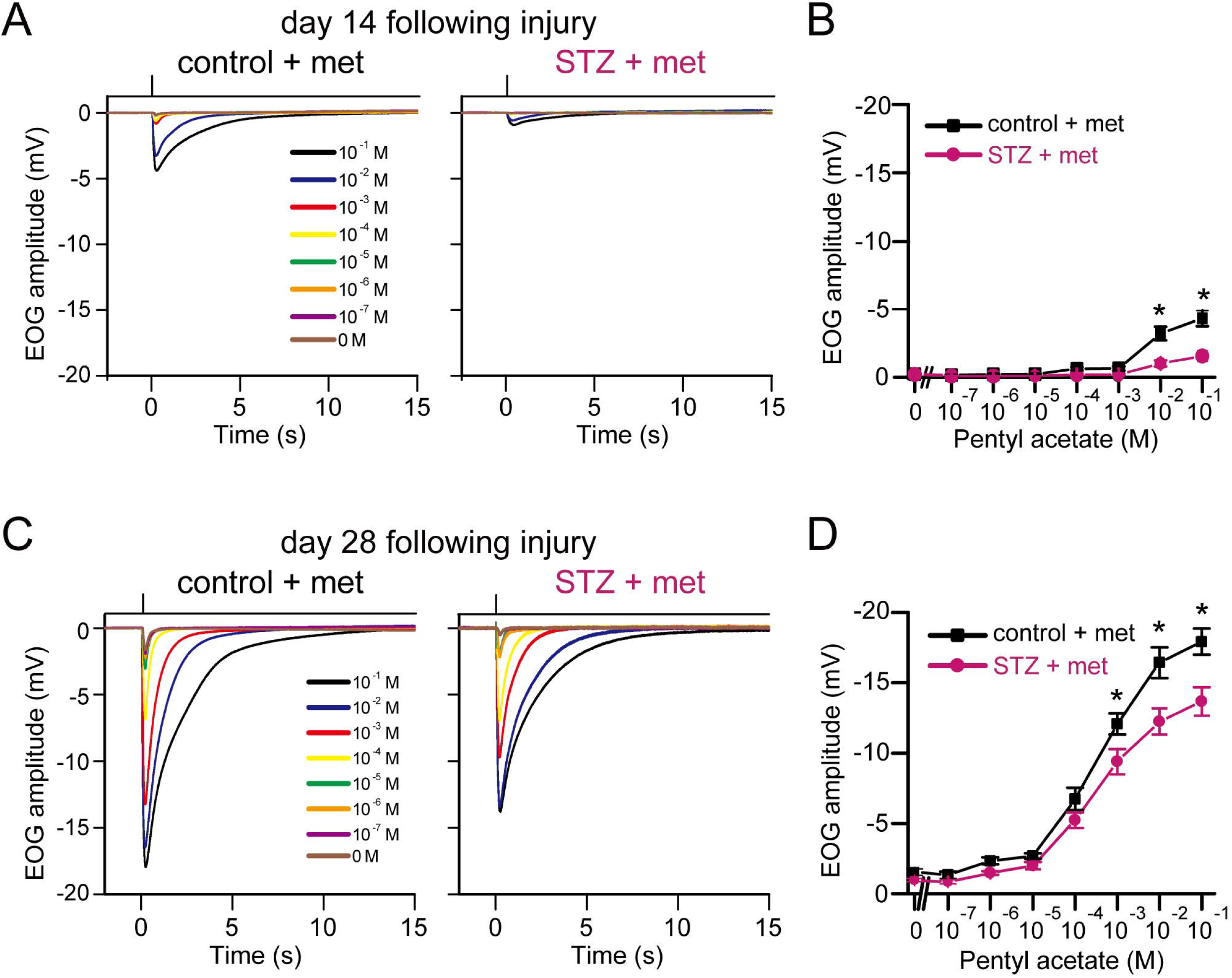
OSNs in STZ mice display reduced odorant-evoked responses. ***A***, Odorant-evoked EOG responses to pentyl acetate at different concentrations in control and STZ mice on day 14 following injury. Similar response kinetics of the EOG were observed in control and STZ mice. ***B***, Comparison of peak amplitudes in EOG recordings between control and STZ mice (*n* = 6 mice/group; * *p* < 0.05, Student’s *t* test) on day 14 following injury. Relative to control mice, STZ mice showed significantly lower EOG amplitudes in response to high concentrations of pentyl acetate (10^−1^ M, *p* = 0.002; 10^−2^ M, *p* = 0.002; 10^−3^ M, *p* = 0.018; Student’s *t* test). Error bars, SEM. ***C***, Odorant-evoked EOG responses to pentyl acetate at different concentrations in control and STZ mice on day 28 following injury. Similar response kinetics of the EOG were observed in control and STZ mice. ***D***, Comparison of peak amplitudes in EOG recordings between control and STZ mice (*n* = 6 mice/group; * *p* < 0.05, Student’s *t* test) on day 28 following injury. Relative to control mice, STZ mice showed significantly lower EOG amplitudes in response to high concentrations of pentyl acetate (10^−1^ M, *p* = 0.012; 10^−2^ M, *p* = 0.015; 10^−3^ M, *p* = 0.044; Student’s *t* test). Error bars, SEM.

### Reduced axonal projections of new OSNs to glomeruli and decreased glomerular responses to odorants in STZ mice

Newly generated OSNs extend their axons to the glomeruli in the OB and form excitatory synapses on the dendrites of interneurons and OB projection neurons within the glomerular structure. As OMP is expressed throughout the axons and axon terminals of mature OSNs (Mori and Sakano, 2011), we measured the OMP-stained areas within individual glomeruli in control and STZ mice to examine whether decreased insulin signaling affects the axonal projections of newly generated OSNs after OE injury. Fig. 4*A* shows representative images of OMP immunostaining in the medial areas of the OB. The axonal target glomeruli in these areas were preferentially selected because they receive projections from the medial part of the OE, including the nasal septum, which we examined for regeneration. Consistent with the results of OMP-positive OSNs in the OE, the OMP-stained areas in the OB on days 14 and 28 postinjury were significantly smaller in STZ mice than in control mice (control-d14 (*n* = 3 mice) vs. STZ-d14 (*n* = 4 mice), *p* < 0.001; control-d28 (*n* = 5 mice) vs. STZ-d28 (*n* = 4 mice), *p* < 0.001; Mann-Whitney *U* test) (Fig. 4*B*). These results suggest that decreased insulin signaling results in axonal projections from fewer newly generated OSNs.

**Figure 4.**
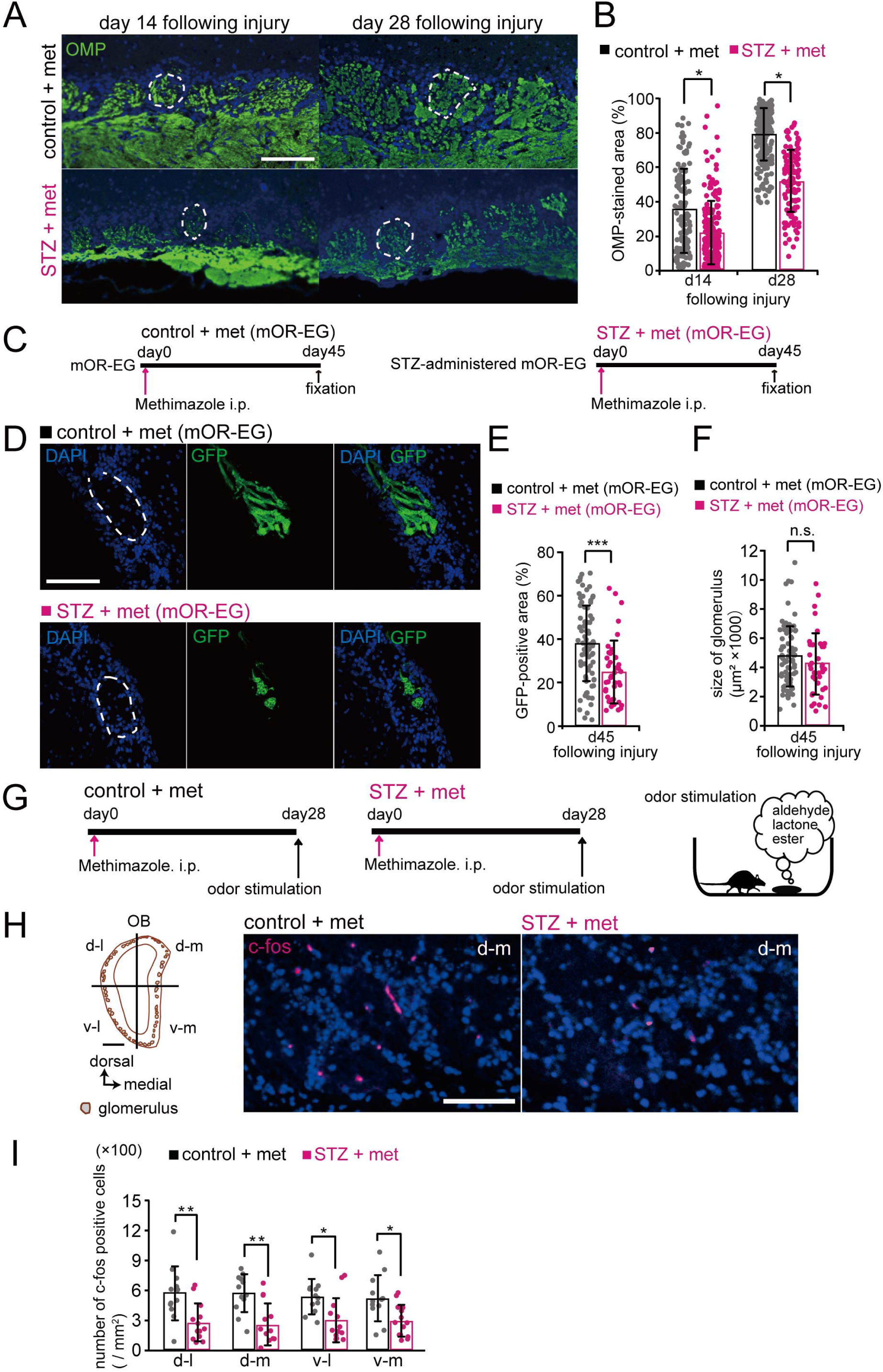
Decreased insulin impairs axonal targeting to glomeruli in the OB. ***A***, Representative coronal sections stained with anti-OMP antibody 14 and 28 days after methimazole-induced injury in control and STZ mice. Each circled area corresponds to a glomerulus. Scale bar, 100 µm. ***B***, Percentages of OMP-stained areas in the medial glomeruli of the OB. The percentages were significantly reduced in STZ mice 14 and 28 days after methimazole-induced injury [d14: 3 mice (control), 4 mice (STZ); d28: 5 mice (control), 4 mice (STZ); * *p* < 0.05, Mann-Whitney *U* test]. ***C***, Time course and experiment design used to test control and STZ mOR-EG-GFP mice. Methimazole was injected i.p., and the tissue was fixed 45 days later. ***D***, Representative sections of GFP-expressing glomeruli in control and STZ mOR-EG-GFP mice 45 days after injury. Dashed circles, glomeruli. Scale bar, 100 µm. ***E***, Percentages of GFP-positive areas within glomeruli. The GFP-labeled area was significantly reduced in STZ-administered mOR-EG-GFP mice (*** *p* < 0.001, Mann-Whitney test). ***F***, Sizes of glomeruli. The mean glomerular sizes were not significantly different between control and STZ mOR-EG-GFP mice (Mann-Whitney test). ***G***, Timeline for odorant-induced c-Fos expression experiment. The fixation and the immunostaining were performed for control and STZ mice 28 days after methimazole-induced injury. Odorants in three categories (aldehydes, lactones, and esters) were applied by placing the odor-containing dish in a cage twice for 1 h with a 10-min interval between placements (right). ***H***, Representative coronal sections of the OB stained with anti-c-Fos antibody. Left, schematic of a coronal OB displaying four quadrants. Right, c-Fos expression of the d-m areas in control and STZ mice. Scale bar, 100 µm. ***I***, Density of c-Fos-positive cells in the glomerular layers in each quadrant of the OB. The densities were significantly smaller in STZ mice than in control mice 28 days after methimazole-induced injury (* *p* < 0.05; ** *p* < 0.01; Mann-Whitney *U* test).

We next investigated whether odorant receptor-specific axonal targeting to specific glomeruli was disturbed in mOR-EG-GFP (as well as I7 and M71) mice (Fig. 4*C*). Very little axonal reinnervation was observed at day 30 (data not shown). Fig. 4*D* shows representative images of GFP-expressing glomeruli from control and STZ-treated mOR-EG-GFP mice 45 days after methimazole injection. Whereas axon terminals of OSNs abundantly spread laterally within a glomerulus from control mice, the terminals were restricted to a limited area of the glomerulus in STZ (mOR-EG-GFP) mice (Fig. 4*D*). Quantitative analyses of GFP-positive areas revealed that the GFP-positive area was significantly smaller in STZ mOR-EG-GFP mice than in control mice at 45 days after injury (control (*n* = 6 mice), STZ (*n* = 4 mice); *p* < 0.001; Mann-Whitney *U* test) (Fig. 4*E*). However, the area of GFP-expressing glomeruli was not significantly different between the control and STZ mOR-EG-GFP mice (*p* = 0.140; Mann-Whitney *U* test) (Fig. 4*F*), suggesting that the reduced GFP-expressing area was a result of impaired axonal targeting. However, incomplete regeneration 45 days after injury was observed in OSNs from both control and STZ I7 and M71 mice (data not shown). Although we cannot rule out the possibility that the time courses for replacing neurons following injury differ for individual odorant receptors, the impaired replacement of axon terminals within glomeruli in STZ (mOR-EG-GFP) mice is consistent with the overall impaired recovery of OMP-positive axon terminals within the OB observed in the STZ-treated C57BL/6 mice.

To determine whether the reduced axonal reinnervation was accompanied by a decrease in the glomerular responses to odorants, we quantified c-Fos (a neural activity marker) induction in cells throughout the OB. At 28 days after methimazole-induced injury, control and STZ mice were exposed to odorants (aldehydes, lactones, and esters) selected for their activation of the dorsal and ventral OB (Fig. 4*G*). A schematic of a coronal section through the OB (Fig. 4*H*, left) displays the four quadrants: ventromedial (v-m), ventrolateral (v-l), dorsomedial (d-m), and dorsolateral (d-l). Representative coronal sections of the d-m area stained by the anti-c-Fos antibody (red) and DAPI (blue) are shown for control and STZ mice in Fig. 4*H* (right). STZ mice had significantly fewer c-Fos-positive cells in each quadrant of the OB than did control mice [control (*n* = 4 mice) vs. STZ (*n* = 5 mice); d-l, *p* = 0.005; d-m, *p* = 0.002; v-l, *p* = 0.007; v-m, *p* = 0.008; Mann-Whitney *U* test; Fig. 4*I*]. Together, these results suggest that the loss of insulin signaling during the recovery from OE injury impairs glomerular olfactory responses because of the incomplete incorporation of new OSNs into olfactory neural circuits.

### STZ mice exhibit impaired detection of the odor of buried food

To determine whether the reduced olfactory signaling in the OE and OB in STZ mice was accompanied by behavioral deficits, mice performed an odor-guided food-seeking paradigm 28 days after OE injury, in which their latency to locate a cookie buried in the bedding was measured (Fig. 5*A,B*). On trial day 1, five of seven STZ mice failed to locate the cookie within the 10 min test time, whereas only one of seven control mice failed this test. On average, STZ mice were significantly slower in locating the cookie (*n* = 7 mice/group; *p* < 0.001, Student’s *t* test) (Fig. 5*C*). The control and STZ mice began to locate the cookie faster on subsequent trial days, but the STZ mice continued to display longer latencies than the control mice (trial day 2, *p* < 0.001; trial day 3, *p* < 0.001, Student’s *t* test) (Fig. 5*C*). On trial day 4, the cookie was not buried and so could be detected visually by the mice. In this trial, STZ mice performed the task as quickly as did the control mice (*p* = 0.463, Student’s *t* test) (Fig. 5*C*). This indicates that no gross motor or motivational deficits account for the observed difference when the cookie was buried. These results suggest that STZ mice have functional deficits in recognizing and/or locating the source of some odors.

**Figure 5.**
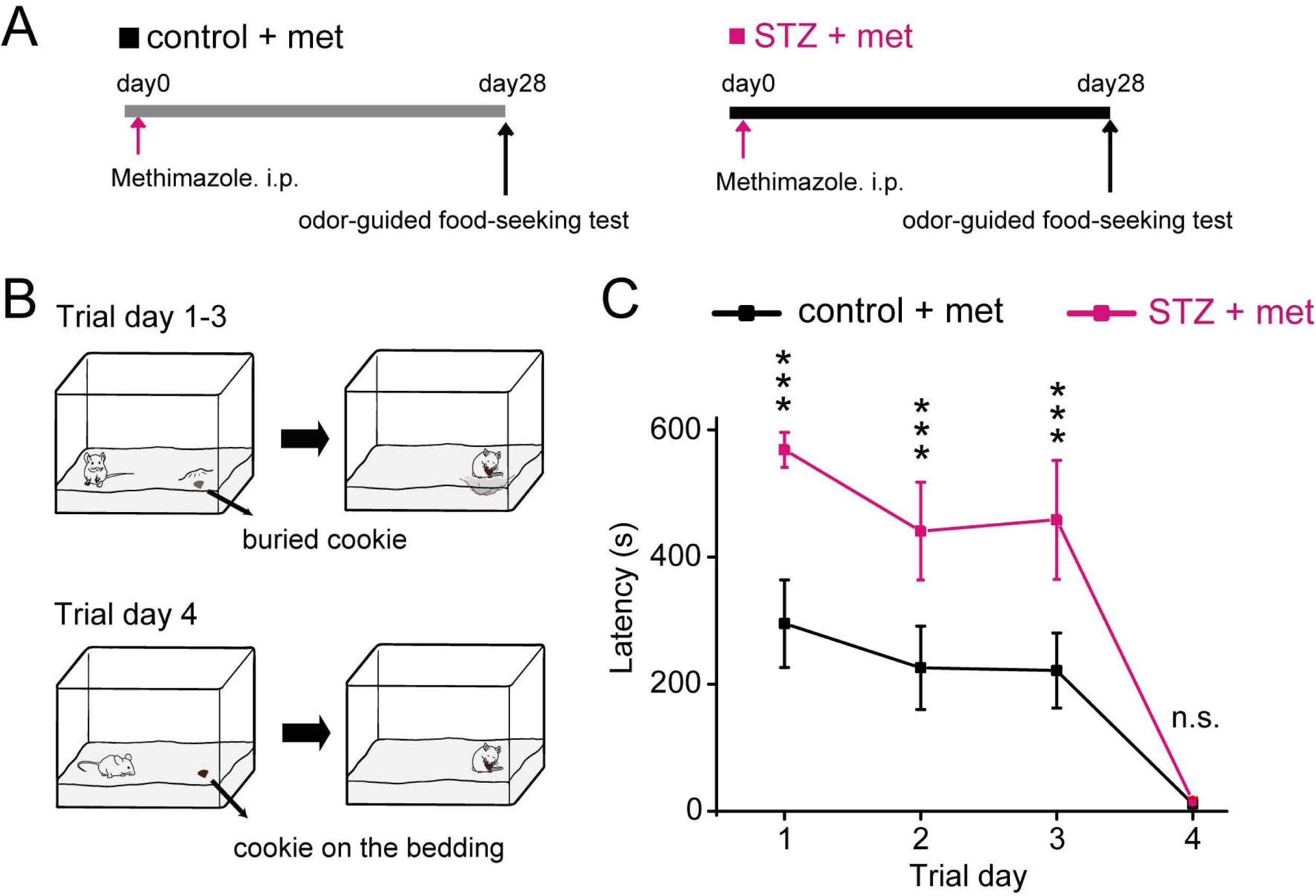
Decreased insulin elicits behavioral deficits in STZ mice 28 days after methimazole-induced injury. ***A***, Time course of the experimental design for the odor-guided food-seeking test. ***B***, Diagram displaying the experimental design of the buried food test. On trial days 1–3, a piece of cookie was buried under the bedding in a randomly selected corner of the mouse cage. On trial day 4, a piece of cookie was placed on top of the bedding to be visible to the mice. ***C***, Latencies to find the cookie on each trial day in control and STZ mice 28 days after methimazole-induced injury (error bars, SEM; *** *p* < 0.001, Student’s *t* test).

### Decreased insulin signaling does not alter proliferation but increases apoptosis in immature neurons

To examine the mechanisms underlying the incomplete replacement of new OSNs after injury, we examined the proliferation and induction of apoptosis of OE cells via immunostaining for Ki67 and activated caspase-3, respectively, 3, 7, 14, and 28 days after injury in control and STZ mice. The representative images of nasal septa in Fig. 6*A* show that the number of Ki67-positive cells began to gradually decrease after the injury (Fig. 6*A*). However, the numbers of Ki67-positive cells did not differ between control and STZ mice at any time point after injury (*n* = 2– 3 mice/group; day 3, *p* = 0.918; day 7, *p* = 0.692; day 14, *p* = 0.224; day 28, *p* = 0.433; Mann-Whitney *U* test) (Fig. 6*B*). By contrast, representative images of activated caspase-3 immunostaining revealed the presence of more apoptotic cells in STZ than in control mice (Fig. 6*C*). Very few activated caspase-3-positive cells were detected in the nasal septa of the OEs from STZ and control mice 7 days after injury (*n* = 3 mice/group; *p* = 0.999; Mann-Whitney *U* test), the numbers of apoptotic cells were significantly higher in STZ mice at the later time points (day 14, *p* < 0.001; day 28, *p* < 0.001; Mann-Whitney *U* test) (Fig. 6*C,D*). The apoptosis of cells in STZ mice was greatest on day 14 (day 7 vs. day 14, *p* < 0.001; day 14 vs. day 28, *p* < 0.001; day 7 vs. day 28, *p* < 0.001; Steel-Dwass test) (Fig. 6*D*). To examine whether these apoptotic cells were mature or immature OSNs, we performed double immunostaining for activated caspase-3 and OMP or GAP43 (Fig. 6*E*). The majority of the activated caspase-3-positive cells were not co-labeled with OMP [91/105 (87%), *n* = 3 mice; white arrowheads in Fig. 6*E*, top] but instead with GAP43 [71/85 (84%), *n* = 3 mice; white arrowheads in Fig. 6*E*, bottom], indicating that nearly all apoptotic cells were immature OSNs. Altogether, these results suggest that the incomplete recovery of OSNs in the OEs of mice with decreased insulin signaling was not a result of reduced proliferation of basal progenitor cells but rather an increase in the apoptosis of OSNs before they reach maturity, which varied between 7 and 28 days after injury.

**Figure 6.**
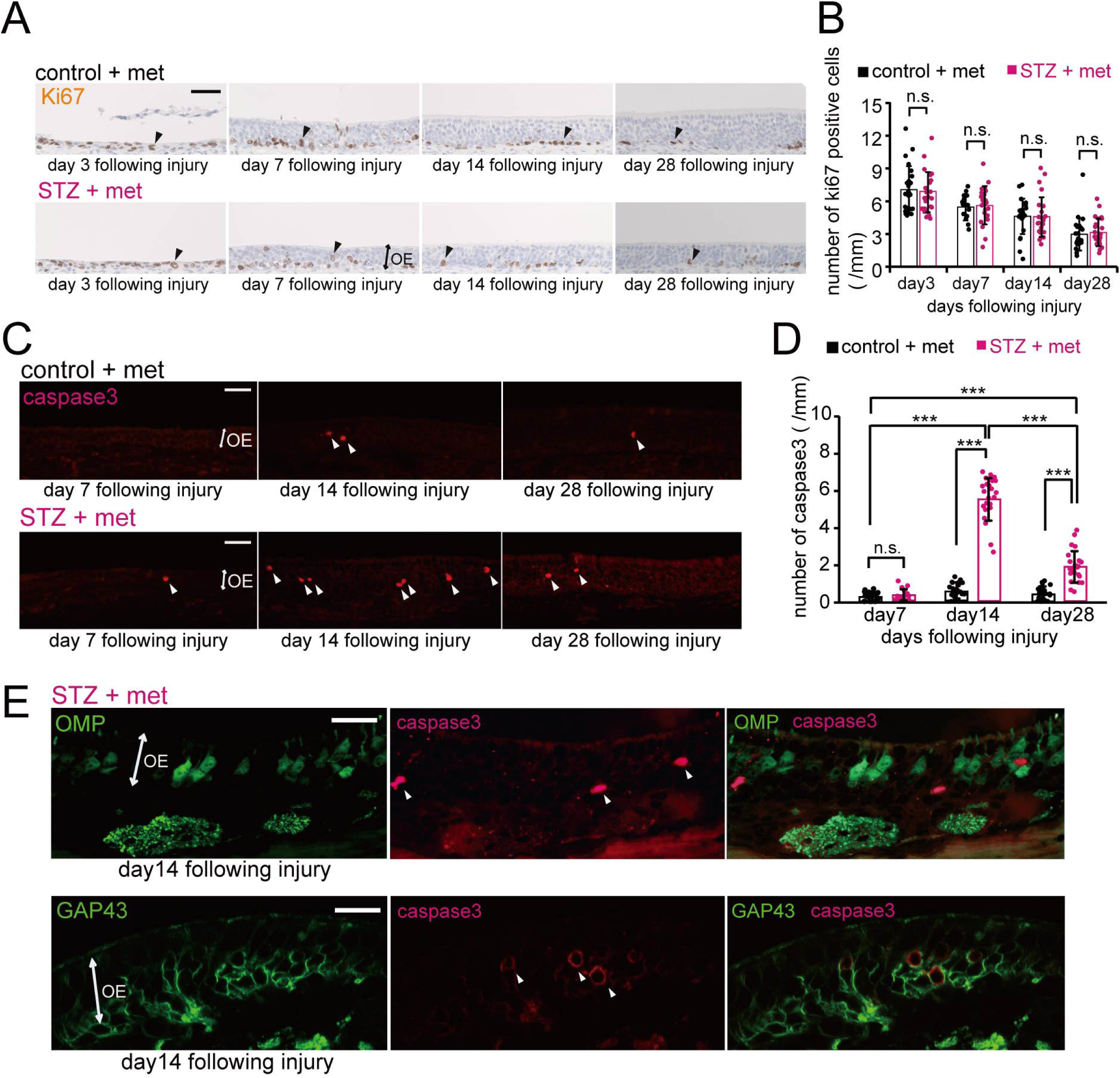
Decreased insulin does not alter OSN proliferation but increases apoptotic cell death in immature neurons. ***A***, Representative coronal sections of nasal septa stained with anti-Ki67 antibody for control and STZ mice 3, 7, 14, and 28 days after methimazole-induced injury. Arrowheads indicate Ki67-positive cells. Scale bar, 50 μm. ***B***, Density of Ki67-positive cells in the nasal septa of control and STZ mice 3, 7, 14, and 28 days after methimazole-induced injury. No significant differences in the numbers of Ki67-positive cells were observed between control and STZ mice [d3: 3 mice (control), 3 mice (STZ); d7: 2 mice (control), 3 mice (STZ); d14: 3 mice (control), 3 mice (STZ); d28: 3 mice (control), 3 mice (STZ); Mann-Whitney *U* test]. ***C***, Representative coronal sections of nasal septa stained with anti-activated caspase-3 antibody for control and STZ mice 7, 14, and 28 days after methimazole-induced injury. Arrowheads indicate activated caspase-3-positive cells. Scale bars, 50 μm. ***D***, Density of activated caspase-3-positive cells in the nasal septa of control and STZ mice 7, 14, and 28 days after methimazole-induced injury. The susceptibility to apoptosis of new OSNs in STZ mice was greatest on day 14 (*n* = 3 mice; ** *p* < 0.01; *** *p* < 0.001; Steel-Dwass test). On days 14 and 28, the total numbers of caspase-3-positive cells in STZ mice were significantly higher than those in control mice (*n* = 3 mice; ** *p* < 0.01; *** *p* < 0.001; Mann-Whitney *U* test). ***E***, Representative coronal sections of nasal septa stained with anti-activated caspase-3 (red) and anti-OMP (green) or anti-GAP43 (green) antibodies 14 days after methimazole-induced injury in STZ mice. The majority of activated caspase-3-positive cells (white arrowheads) were costained not by the anti-OMP but by the anti-GAP43 antibody. Scale bar, 20 µm.

### Period of susceptibility for insulin signal-dependent survival or death of new OSNs

We previously reported that new OSNs have a specific time window for sensory input-dependent survival or death and are more susceptible to apoptosis when they become OMP-positive mature cells (Kikuta et al., 2015). We hypothesized that this enhanced susceptibility would also occur with decreased insulin signaling. To identify a period during which new OSNs may be highly dependent on insulin signaling, we investigated the replacement of new OSNs in response to insulin administration at different times after injury. We focused on the first 14 days after injury, because it was during this period that we observed the emergence of OMP-positive neurons.

In this experiment, insulin was administered to STZ mice via i.p. injections throughout the 14-day postinjury period or during the first or second half of this period (Fig. 7*A*). Fig. 7*B* shows representative coronal sections of the nasal septa from control and STZ mice 14 days after injury. The OEs of mice that received insulin on days 1–6 were significantly thinner, with fewer OSNs and OMP-positive cells, than the OEs of mice that received insulin on days 1–13 and days 8–13 (*n* = 3–4 mice/group; OE thickness, d1–13 vs. d1–6, *p* < 0.001; d1–6 vs. d8–13, *p* < 0.001; d1–13 vs. d8–13, *p* = 0.285; number of OSNs, d1–13 vs. d1–6, *p* < 0.001; d1–6 vs. d8–13, *p* < 0.001; d1–13 vs. d8–13, *p* = 0.367; number of OMP-positive cells, d1–13 vs. d1–6, *p* < 0.001; d1–6 vs. d8–13, *p* < 0.001; d1–13 vs. d8–13, *p* = 0.499; Mann-Whitney *U* test) (Fig. 7*C–E*). By contrast, there were no differences in these parameters between mice receiving insulin on days 8–13 and those receiving insulin for the entire period (*n* = 3–4 mice/group; OE thickness, *p* = 0.323; number of OSNs, *p* = 0.402; number of OMP-positive cells, *p* = 0.499; Mann-Whitney *U* test) (Fig. 7*C–E*). These results suggest that newly generated OSNs are highly dependent on insulin signaling 8–13 days post injury.

**Figure 7.**
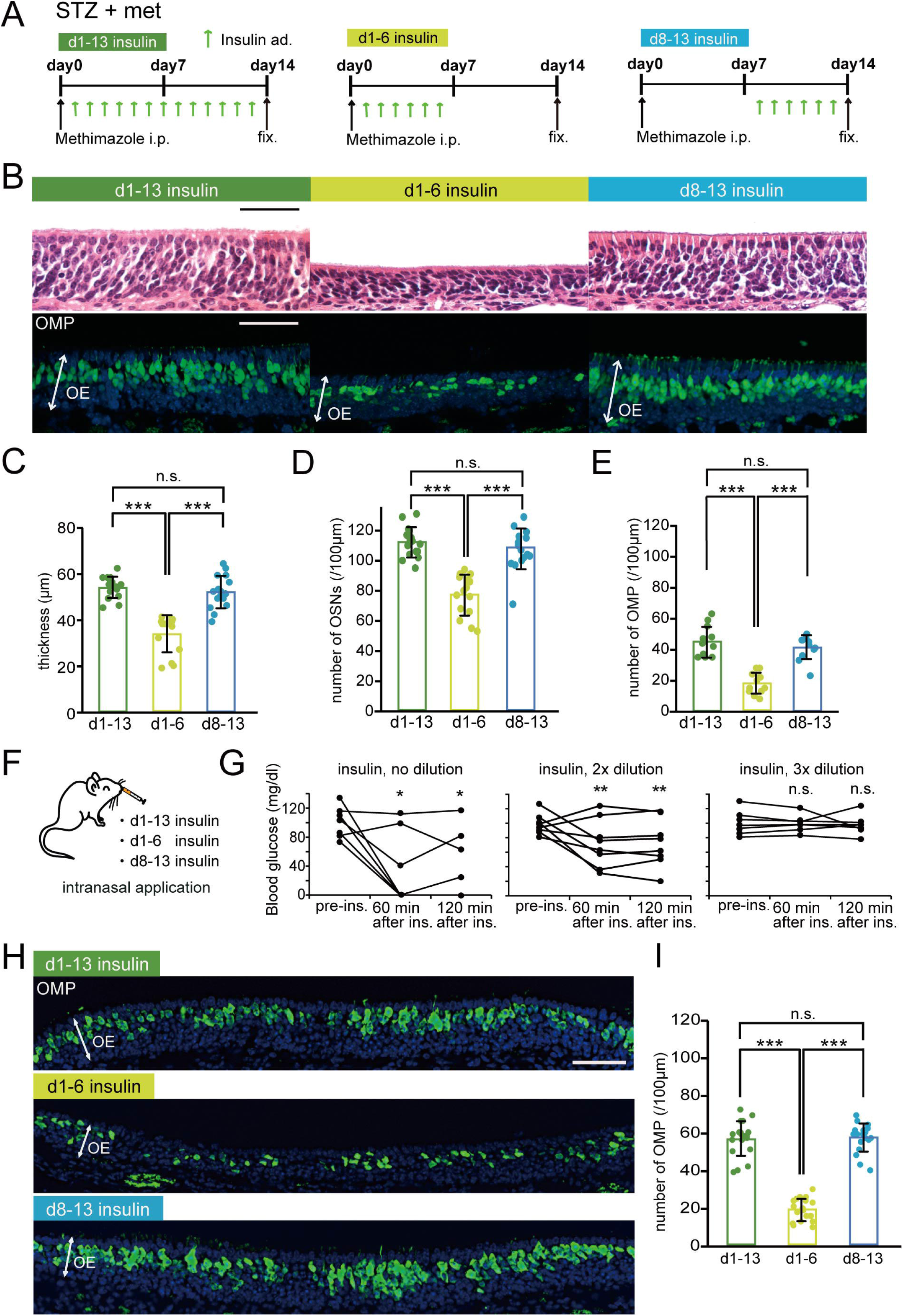
Newly generated OSNs require insulin during their maturation after day 7 post injury. ***A***, Time course of the experimental design. Mice in three experimental groups were administered insulin i.p. at different times following methimazole-induced injury. ***B***, Representative coronal sections of the nasal septa stained with hematoxylin (upper) or anti-OMP antibody (lower) for mice administered insulin on d1–13, d1–6, or d8–13. Scale bars, 50 µm. ***C–E***, OE thicknesses (***C***), density of OSNs (***D***), and density of OMP-positive cells (***E***) in the three groups [OE thickness: 3 mice (d1–13), 3 mice (d1–6), 3 mice (d8–13); number of OSNs: 3 mice (d1–13), 3 mice (d1–6), 3 mice (d8–13); number of OMPs: 4 mice (d1–13), 3 mice (d1–6), 3 mice (d8–13); * *p* < 0.05; *** *p* < 0.001; Mann-Whitney *U* test]. ***F***, Diagram of intranasal insulin administration. Insulin was applied to the nasal cavities of STZ mice after methimazole-induced injury. These applications were performed according to the protocol used for the i.p. insulin administration shown in ***A***. ***G***, Effects of the intranasal insulin application on blood glucose levels. Shown are blood glucose levels before (pre-ad) and 60 and 120 min after intranasal administration (after ad) of insulin at three concentrations (* *p* < 0.05; ** *p* < 0.01; Steel test). ***H***, Representative coronal sections of nasal septa stained with anti-OMP antibody (green) 14 days after methimazole-induced injury in STZ mice. Scale bar, 50 µm. ***I***, Density of OMP-positive cells 14 days after methimazole-induced injury in the three intranasal insulin administration groups (*n* = 3 mice; *** *p* < 0.001; Mann-Whitney *U* test).

Blood glucose levels are inevitably altered by i.p. insulin administration, which may affect the regeneration of the OE after injury via increased superoxide production (Giacco and Brownlee, 2010). Thus, it remains unclear whether the impaired OE regeneration in STZ mice was influenced by the loss of insulin signaling in the OE or by the high blood glucose levels caused by hypoinsulinemia. To investigate this, we examined the effects of intranasally applied insulin on OE regeneration in STZ mice (Fig. 7*F*). We first determined which insulin concentration did not affect blood glucose levels. Fig. 7*G* shows blood glucose levels before (pre-ad) and at 60 and 120 min after intranasal administration of insulin at three concentrations. An insulin solution diluted 3× with saline did not alter blood glucose levels after 60 or 120 min (*n* = 7 mice; pre-ad vs. 60 min, *p* = 0.973; pre-ad vs. 120 min, *p* = 0.928; Steel test) (Fig. 7*G*), whereas the higher concentrations significantly reduced the blood glucose levels (no dilution, *n* = 7 mice: pre-ad vs. 60 min, *p* = 0.044; pre-ad vs. 120 min, *p* = 0.046; 2× dilution, *n* = 8 mice: pre-ad vs. 60 min, *p* = 0.002; pre-ad vs. 120 min, *p* = 0.003; Steel test) (Fig. 7*G*). Thus, we used the 3× diluted insulin solution to examine the effect of insulin on OE regeneration in the absence of altered blood glucose levels. Fig. 7*H* shows representative coronal sections of nasal septa from STZ mice for each treatment regime 14 days after injury. Consistent with the results from i.p. insulin administration, the number of OMP-positive OSNs was significantly lower only in the group receiving intranasal insulin only on days 1–6 (d1–13 vs. d1–6, *p* < 0.001; d1–6 vs. d8–13, *p* < 0.001; d1–13 vs. d8–13, *p* = 0.929; Mann-Whitney *U* test) (Fig. 7*I*). These results indicate that the absence of insulin signaling 8–13 days after injury impairs the recovery of the OE. Thus, insulin signaling during this period is involved in the regeneration of the OE.

### Upregulation of insulin signaling during the specific period is required for OSN maturation

**Fig. 7** shows that insulin-dependency is increased on days 8–13 post-injury, implying upregulation of *Insr* expression during this period. To examine whether the level of *Insr* expression is changed between uninjured OE and the OE at day 14 following injury, an RNAscope assay was carried out using an *Insr* probe. Fig. 8*A,B* show representative images of the uninjured OE and OB and the OE and OB at day 14 after injury, with the *Insr* mRNA signal appearing as small brown dots. Similar to *in situ* hybridization results in Fig. 1*A,B*, the RNAscope signals are observed sparsely in the layer of supporting cells and OSNs of the uninjured OE (Fig. 8*A* left panel). Overall, the signal intensity is relatively weak. However, Fig. 8*A* (right panel, day 14 following injury) shows a much higher number of dots both in supporting cell and OSN layers, while the mRNA signal intensity appears similar between the uninjured OB and the OB at day 14 following injury (Fig. 8*B*). No signals were detected by the negative control probe B. subtilis dihydrodipicolinate reductase (DapB) in the OE of control mice (data not shown). These results suggest that *Insr* expression is highly upregulated in the OE at day 14 post-injury, when the delay of OE regeneration after injury had already begun in STZ-mice after day 7 post-injury (Fig. 2*B–F*, Fig. 3*A,B*, Fig. 4*A,B*, Fig. 6*C–E*).

Given these results, we next applied intranasally the insulin receptor antagonist, S961 to assess the effect of blocking insulin signaling on OE regeneration following injury. In this experiment, S961 was applied on days 8–13 post-injury twice daily and intranasal PBS application was performed similarly with control mice (Fig. 8*C*). To compare the effect of blocking insulin signaling within a mouse, S961 or PBS was unilaterally applied (Fig. 8*D*). We also monitored the effect of intranasal S961 application (0.5 µg/µl, 10µl) on blood glucose level. Fig. 8*E* shows the blood glucose values before and at 120 min after intranasal S961 application. No obvious increase of blood glucose by intranasal S961 application was observed (*n* = 4 mice; pre-S961 vs. 120 min) (Fig. 8*E*), suggesting that this treatment did not alter systemic glucose levels. Fig. 8*F* shows the representative images of coronal sections of the nasal septum on day 14 post-injury in control mice treated with PBS or S961. PBS application did not cause any significant decreases of the OE thickness and the number of OSNs and OMP-positive OSNs compared with those in the OE of the control side (*n* = 3 mice/group; OE thickness, *p* = 0.863; number of OSNs, *p* = 0.436; number of OMP-positive cells, *p* = 0.605; Mann-Whitney *U* test) (Fig. 8*G–I*). By contrast, supporting the results from intranasal insulin application (Fig. 7*H,I*), the OE thickness and the number of OSNs and OMP-positive OSNs were significantly smaller in the OE that received S961 compared with those in the OE of the control sides (*n* = 3 mice/group; OE thickness, PBS-C vs. S961, *p* < 0.001; S961 vs. S961-C, *p* < 0.001; number of OSNs, PBS-C vs. S961, *p* < 0.001; S961 vs. S961-C, *p* < 0.001; number of OMP-positive cells, PBS-C vs. S961, *p* < 0.001; S961 vs. S961-C, *p* < 0.001; Mann-Whitney *U* test) (Fig. 8*G–I*). Taken together, these results suggest that insulin signaling via the insulin receptor is required for OSN maturation at days 8–13 post-injury, when *Insr* expression is highly upregulated in the OE.

**Figure 8.**
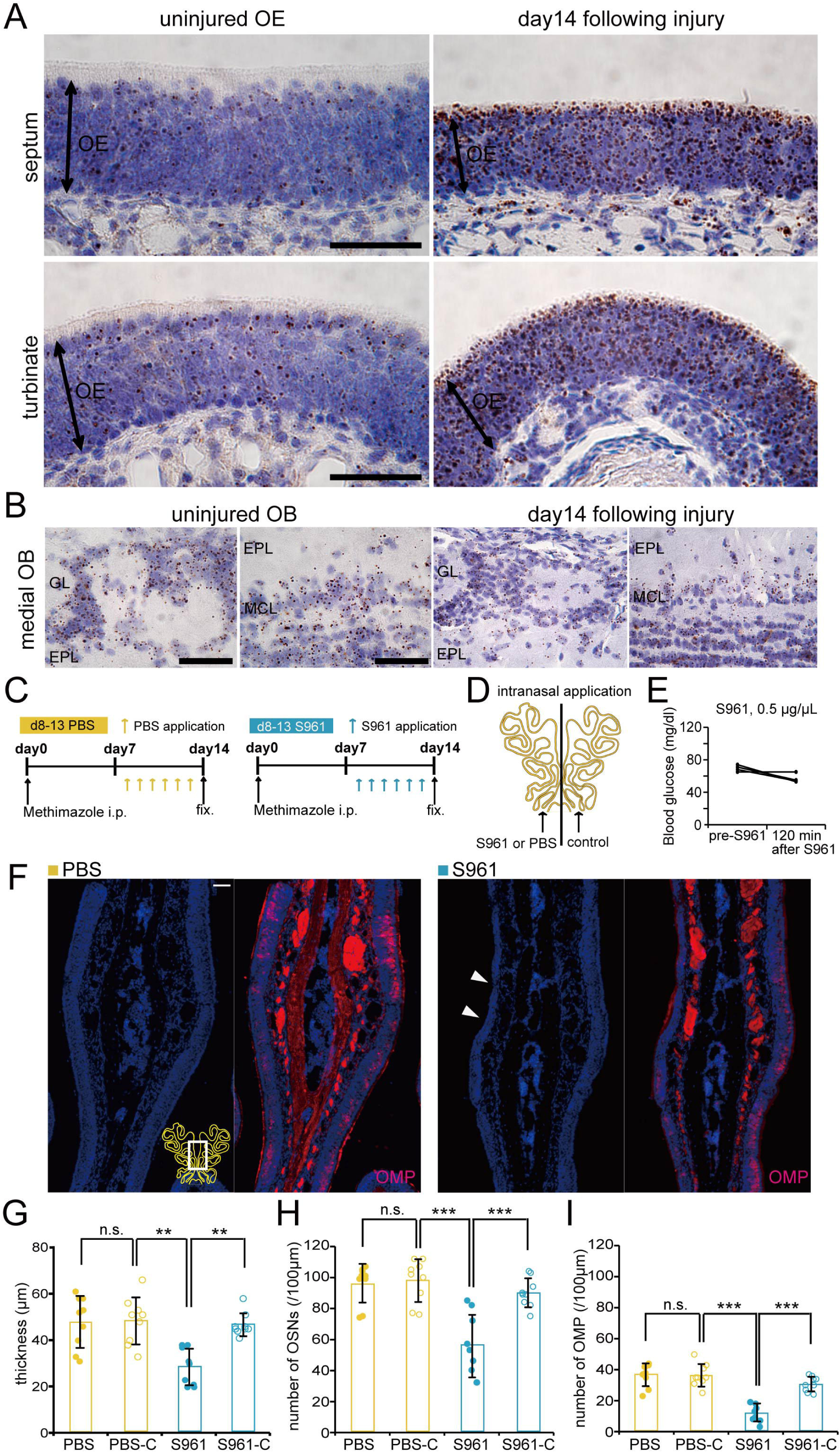
Upregulation of insulin receptor-mediated during day 8–13 is required for maturation of OSNs. ***A, B***, *In situ* hybridization by RNAscope assay of *Insr* from septal and turbinate coronal sections of the uninjured OE and the OE on day 14 following injury (***A***) and medial part of sections of the uninjured OB and the OB on day 14 following injury (***B***) in non-diabetic 10-wk-old mice. Signals were sparsely detected especially in the layer of supporting cells and OSNs of the uninjured OE (*n* = 3 mice) (left panels in ***A***). Strong signals, suggesting upregulation of *Insr* expression were detected in the OE on day 14 following injury (*n* = 3 mice) (right panels in ***A***). In the OB, signals were detected in the glomerular layer (GL), external plexiform layer (EPL), and mitral cell layer (MCL) (***B***). Signal intensity appears similar between the uninjured OB (*n* = 3 mice) (left panels in ***B***) and the OB at day 14 following injury (*n* = 3 mice) (right panels in ***B***). Scale bars, 50 µm. ***C***, Time course of the experimental design. Mice in two experimental groups were administered PBS or S961, an insulin receptor antagonist during day 8–13 following methimazole-induced injury. ***D***, Diagram of unilateral intranasal application. PBS or S961 was applied to a side of the nasal cavities of non-diabetic control mice after methimazole-induced injury according to the protocol shown in ***C***. ***E***, Effects of the intranasal S961 application (0.5 µg/µl, 10 µl) on blood glucose levels before (pre-S961) and 120 min after intranasal administration (after S961) of insulin. ***F***, Representative coronal sections of the nasal septa stained with DAPI and anti-OMP antibody (red) for control mice intranasally administered PBS or S961 on d8–13. Scale bars, 50 µm. ***G–I***, OE thicknesses (***G***), density of OSNs (***H***), and density of OMP-positive cells (***I***) in the three groups [OE thickness: 3 mice (PBS), 3 mice (S961); number of OSNs: 3 mice (PBS), 3 mice (S961); number of OMPs: 3 mice (PBS), 3 mice (S961); * *p* < 0.05; ** *p* < 0.01; *** *p* < 0.001; Mann-Whitney *U* test]. PBS-C, PBS-Control; S961-C, S961-Control.

### Insulin signal promotes the OE regeneration following injury when *Insr* expression is highly upregulated

If insulin signaling is important for the maturation and survival of newly-generated OSNs following injury, when *Insr* expression is highly upregulated, an insulin-enriched environment could facilitate their maturation even in non-diabetic mice. To test this hypothesis, we first examined the effect of intranasal insulin application on OSN regeneration after injury (in non-diabetic mice). Insulin was applied to a single naris on days 1–6 and days 1–13 postinjury three times daily to compare the effect of insulin signaling between the two sides of the nose (Fig. 9*A*). Representative images of the nasal septum stained with anti-OMP antibody for both groups are shown in Fig. 9*B*. In the group subjected to insulin application 1 – 6 days after injury, OMP-positive cells were not identified at this stage in both the application and contralateral sides. At 14 days after injury, newly generated OSNs began to mature, but intranasal application during 1–13 days after injury induced a significant increase of OMP-positive cells on the application side compared with the contralateral side (*n* = 4 mice; number of OMP-positive cells, *p* < 0.001; Mann-Whitney U test) (Fig. 9*C*). These results indicate that insulin application 1–13 days after injury increases and further facilitates the maturation of newly generated OSNs even in non-diabetic mice, leading to faster functional recovery of the OE. To determine whether insulin signaling during the period of high *Insr* expression would be critical for the promotion of functional recovery of the OE following injury, we examined the effect of insulin application on OSN regeneration in an early period (d1–6 insulin after injury) and a late period (d8–13 insulin after injury). The details of this experiment are described in Fig. 9*D*. The timing of the insulin application differed between the two groups such that one group of mice received insulin on 1–6 days after injury (Fig. 9*D*, left) and the other group received the insulin application on 8–13 days after injury (Fig. 9*D*, right). Representative coronal sections of the nasal septum and concha bullosa for both groups are shown in Fig. 9*E*.

In the group subjected to insulin application 1–6 days after injury, the number of OMP-positive cells for both sides did not differ (*n* = 4 mice; number of OMP-positive cells, *p* = 0.984; Mann-Whitney U test) (Fig. 9*F*). By contrast, insulin application during days 8–13 caused a significant increase of the OMP-positive cells on the application side compared with those on the contralateral side (*n* = 4 mice; number of OMP-positive cells, *p* < 0.01; Mann-Whitney U test) (Fig. 9*F*). Furthermore insulin application during this period caused a significant increase of the OMP-positive cells on the application side compared with those on the application side during days 1-6 after injury (*p* < 0.001; Mann-Whitney U test) (Fig. 9*F*). These results indicate that insulin signaling during the period when *Insr* is highly expressed is a primary factor to facilitate the incorporation of newly generated OSNs into neural circuits.

**Figure 9.**
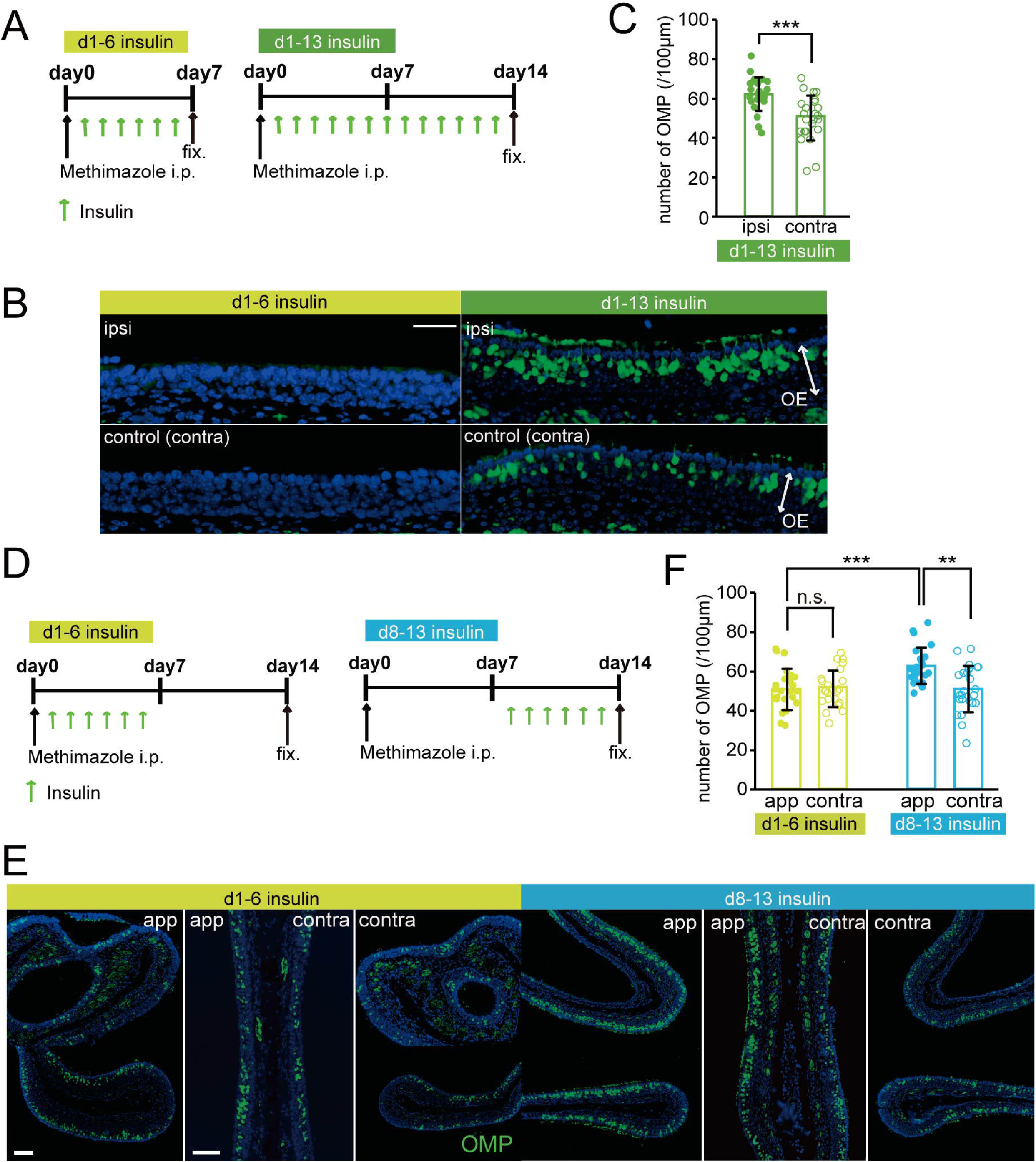
Insulin enhances the replacement of newly generated OSNs in control mice. ***A***, Time course of the experimental design following methimazole and insulin application. Mice in two experimental groups received unilateral intranasal insulin application at different periods after methimazole-induced injury. Both groups received methimazole intraperitoneally (i.p.) on day 0 and intranasal insulin starting on day 1. Mice were perfused with fixative on day 7 (d1-6 insulin, left) or day 14 (d1-13 insulin, right) after the methimazole-induced injury. ***B***, Representative images of the nasal septum stained with anti-olfactory marker protein (OMP) (green) antibody in the two groups. Scale bar, 50 µm. ***C***, Number of olfactory marker protein (OMP)-positive cells in mice subjected to insulin administration 1–13 days after the injury. The number of OMP-positive cells were significantly increased on the insulin application side compared with the contralateral side (\*\*\**p* < 0.001; Mann–Whitney test). ***D***, Time course of the experimental design of methimazole and insulin application. Mice in two experimental groups received unilateral insulin application at different times after methimazole-induced injury. Both groups received methimazole intraperitoneally on day 0. Insulin was unilaterally applied on days 1–6 (left, d1-6 insulin) and days 8–13 (right, d8-13 insulin) post-injury three times daily. Mice were perfused with fixative on day 14 after the methimazole-induced injury. ***E***, Representative images of the olfactory nasal septum and concha bullosa stained with anti-olfactory marker protein (OMP) (green) antibody in the two groups. App, application side; contra, contralateral side. Scale bars, 50 µm. ***F***, Number of olfactory marker protein (OMP)-positive cells in mice subjected to insulin application 1–6 days and 8–13 days after injury. The number of OMP-positive cells on the application side was significantly higher during the insulin application 8-13 days post-injury than that on the contralateral side (\*\*\**p* < 0.001; Mann–Whitney test), whereas no significant difference was observed between the application and contralateral sides 1-6 days post-injury (n.s., not significant; Mann-Whitney test).

## Discussion

This study demonstrates that the insulin receptor is required for regeneration following injury and that its expression in the OE is dynamically regulated depending on the physiological or regenerative condition of the OE. Overall moderate and locally restricted signals in the uninjured OE for *Insr* (Figs 1*A,B*, 8*A*) hint at local regeneration as part of the normal turnover of cells in the OE. And high expression levels of *Insr* in the OE on day 14 after the total degeneration of OSNs and supporting cells induced by methimazole support the requirement of the insulin receptor in the regeneration process (Fig. 8*A*). Intranasal insulin application directly to the OE improves the maturation and survival of new neurons after methimazole-induced injury (Fig. 7*H,I*, Fig. 8*F–I*, Fig. 9). Insulin is a growth-stimulating hormone produced by pancreatic beta cells (Baura et al., 1993; Banks, 2004; Laron, 2009; Fernandez and Torres-Aleman, 2012) and perhaps also in the brain (Gray et al., 2014). Its effects parallel those of insulin-like growth factors IGF-1 and IGF-2 because of their structural similarities and the high degree of homology between the insulin and IGF receptors. However, insulin binds to the insulin receptor with a 100–1,000-fold higher affinity than that for other IGF receptors (Fernandez and Torres-Aleman, 2012). In contrast to insulin binding and insulin receptor levels in many peripheral tissues, those in the brain are not upregulated in experimental diabetes mellitus (Pacold and Blackard, 1979; Sechi et al., 1992; Pezzino et al., 1996). After injury to the OE in non-diabetic mice, we found that *Insr* expression was highly upregulated in the whole OE layer (Fig. 8*A*). Thus, a decreased level of circulating insulin and diminished insulin likely impair the regeneration of the OE.

Under our control conditions (i.e., without OE injury), the OE contained primarily mature OSNs. An STZ-induced decrease in insulin did not affect the structure and odorant-induced responses of the OE (Figs 1, 10*A*). Because *Insr* expression is moderate especially in mature OSNs of the uninjured OE (Figs 1*A,B*, 8*A*) (Saraiva et al., 2015), the dependency on insulin may not be large, resulting in few effects on the OE in STZ mice. After methimazole-induced injury, however, we observed the generation of new OSNs and their gradual incorporation into neural circuits (Figs 2*B,E*, 4*A,D*, 10*B*). The decreased insulin in STZ mice significantly affected the regeneration process 8 – 13 days after the injury. Specifically, hypoinsulinemia reduced the thickness of the OE, the number of OSNs, and the number of OMP-positive cells (Figs 2*B–F*, 10*B*). Consistent with these findings, we observed increased apoptosis but unchanged proliferation of immature OSNs in STZ mice during this period (Fig. 6). The nasally-applied insulin during the regeneration process compensated for and rescued an incomplete repair of the OE in STZ mice (Fig. 7) and promoted the OE repair even in control, non-diabetic mice (Fig. 9), while the block of insulin receptor-mediated signaling impaired the OE repair in non-diabetic mice (Fig. 8). These results indicate that the survival of new neurons requires insulin and the insulin receptor, and their survival strongly influences the regeneration of the OE.

**Figure 10.**
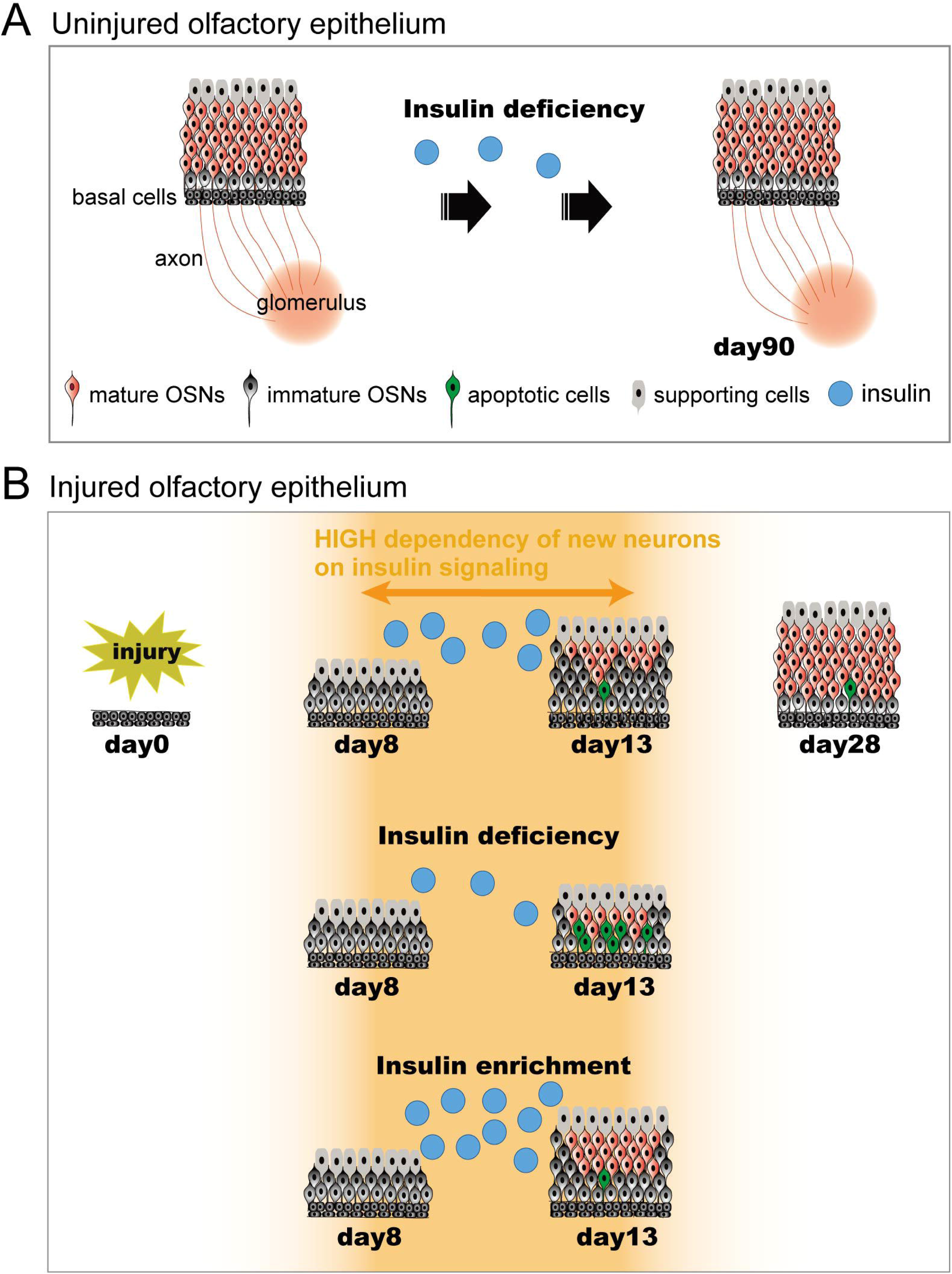
Diagram for insulin-dependent replacement of newly generated OSNs following injury. ***A***, Time course of tissue structure in uninjured OE. Even under insulin-deficient situations, no structural changes in the uninjured OE occur during the 90-day period. ***B***, Time course of the repair process following injury. The dependence of the newly generated OSNs on insulin increased between 8 and 13 days following injury. Under normal insulin levels, newly generated OSNs regenerate and mature during days 8-13 following injury. The number of mature OSNs gradually increases, and tissue repair is completed 28 days following injury. However, under insulin-deficient situations during days 8-13 following injury, newly generated OSNs are highly susceptible to apoptosis, resulting in incomplete recovery of the OE with fewer mature OSNs. Under insulin-enriched situations during the same period, facilitation of the OE repair could occur.

### Possible mechanisms for incomplete replacement of new OSNs in the STZ mice

Insulin in the brain acts as a neuromodulator influencing synaptic plasticity, dendritic outgrowth, and neurotransmitter release, while also promoting neuronal survival and proliferation (Bruning et al., 2000; Plum et al., 2005; Mielke et al., 2006; Valenciano et al., 2006). Thus, insulin is likely strongly involved in the maintenance of neural circuit functions. Activation of the insulin receptor involves intracellular downstream cascades, including phosphoinositide 3-kinase–AKT–forkhead box protein O and RAS-MAPK pathways. These pathways modulate gene transcription and activate myriads of downstream kinase-phosphatase branches that ultimately affect key cellular processes, such as protein synthesis, autophagy, apoptosis, and resistance to oxidative stress (Fernandez and Torres-Aleman, 2012). Furthermore, STZ-treated rats exhibit reduced expression of glycogen synthase kinase 3β in the brain (Jing et al., 2017), which activates the Wnt signaling pathway that reduces synaptic GABA and glutamate transporters important for the maintenance of synapses (Kuwabara et al., 2009). Therefore, decreased insulin may promote apoptosis and Wnt signaling. The suppression of cell death of immature neurons and adequate synaptic transmission at the axon terminals driven by insulin may be key factors in the recovery of the OE and rewiring in the OB.

Our observation of enhanced caspase activation in new neurons of STZ mice might reflect the decreased neuronal activity under reduced insulin. Neuronal activity, which is characterized by the generation of action potentials, leads to increased cytoplasmic cAMP levels (Fiske and Brunjes, 2001; Watt et al., 2004; Turrigiano, 2012; Francois et al., 2013) and subsequent phosphoinositide 3-kinase–AKT signaling (Balazs et al., 1988; Barger, 1999; Watt et al., 2004; Hyman and Yuan, 2012). These signals inhibit the intrinsic apoptosis machinery, thereby preventing cellular suicide (Raff, 1992; Burek and Oppenheim, 1996; Frade et al., 1996; Nunez and del Peso, 1998; Song and Poo, 1999; Vaux and Korsmeyer, 1999; Duarte et al., 2012). Patch-clamp recordings in rats have shown that insulin regulates the spontaneous firing of action potentials in olfactory neurons (Savigner et al., 2009). Furthermore, neuronal activity induces synapse maturation by promoting the incorporation of NMDA receptors containing the subunit 2A and the recruitment of AMPA receptors to the postsynaptic site to activate silent synapses and to increase the strength of synaptic transmission (Isaac et al., 1995; Wu et al., 1996; Li and Sheng, 2003). Accordingly, the loss of insulin may result in reduced activity in OSNs and a failure to establish stabilized synaptic connections with target OB neurons, thereby ultimately contributing to the induction of apoptosis. Importantly, *Insr* is highly expressed in the OB (Fernandez and Torres-Aleman, 2012) and OSN apoptosis occurs around when OMP expression begins and when OSNs are born following OB ablation (Schwob et al., 1992; Coleman JH et al., 2019). Thus, insulin in the OB may also contribute to maturation of OSNs.

### Time-dependent insulin in newly generated OSNs

We observed that insulin deficiency and blocking insulin receptors during days 8–13 after injury reduced the incorporation of new OSNs. During the period, both increased apoptotic cell death of immature OSNs and *Insr* upregulation were observed. Furthermore, nasal insulin application during the same period promoted the maturation of OSNs in control mice. These results indicate that the susceptibility of new OSNs to apoptosis varies according to their maturation stage and that new OSNs have a critical stage (8 – 13 days following injury) during which they are highly dependent on insulin for maturation. This stage corresponds to the growth of axons into the OB and subsequent expression of VGluT2, for the vesicular release of glutamate (Miragall and Monti Graziadei, 1982; Rodriguez-Gil et al., 2015; Liberia T et al., 2019). Insulin also regulates neurite outgrowth in cultured neurons and spine density (Govind et al., 2001; Choi et al., 2005), promotes the surface expression of recombinant NMDA receptors on Xenopus oocytes (Skeberdis et al., 2001), and accelerates the insertion of GluR1-containing AMPA receptors into the membranes of cultured hippocampal neurons (Passafaro et al., 2001). Conversely, an attenuation of insulin signaling reduces AMPA mEPSC frequency and synaptic contacts onto tectal neurons (Chiu and Cline, 2010), leading to functionally incomplete synaptogenesis. Therefore, we speculate that insulin receptor-mediated signaling supports the growth of axons of newly generated OSNs to the target sites and the development of synapses, contributing to their incorporation into and maintenance within the olfactory neural circuits. Under conditions of low (or no) insulin receptor-mediated signaling, the axons of new neurons failed to reach postsynaptic sites and establish appropriate connections with OB neurons, resulting in the activation of apoptotic cascades that would otherwise be prevented with successful and functional integration into the circuit. Under conditions of sufficient insulin levels, the axons of new OSNs effectively reach the OB, resulting in the promotion of OSN maturation. The axon development and formation of functional synapses with OB neurons might be the insulin-dependent processes that explain the insulin-sensitive period in new neurons 8 – 13 days after the injury.

From a clinical perspective, olfactory damage, such as from traumatic injury, viral infections, and rhinosinusitis, diminishes an individual’s quality of life. In some cases, functional recovery from damage is incomplete despite continuous OSN regeneration (Schwob, 2002). Potential therapeutic options that help achieve complete tissue regrowth and functional recovery are not fully established. If insulin is important for the maturation and survival of new OSNs in the OE, intranasal administration represents a potential therapeutic option for olfactory dysfunction. The results from this study indicate that application between 8 - 13 days after injury would be the most effective to promote OE recovery. An improved understanding of the molecular and cellular mechanisms involved in the insulin-dependent period of OE regeneration should facilitate the effective utilization of insulin administration for the recovery of olfactory functions.

## Acknowledgments

This study was supported by a grant-in-aid for scientific research (C) of the Japan Society for Promotion of Science grant 17K11354 (Shu Kikuta), Takeda Science Foundation grant (Shu Kikuta) and a National Institutes of Health grant R01DC016647 (Johannes Reisert). Immunohistochemistry and confocal microscopy were, in part, performed at the Monell Histology and Cellular Localization Core which is supported, in part, by funding from the NIH-NIDCD Core Grant 1P30DC011735-01 and also from grant G20OD020296 (infrastructure improvement at the Monell Chemical Senses Center). We thank Ms. Emi Usukura at the University of Tokyo for technical assistance. We also thank Prof. Kensaku Mori (University of Tokyo) for critical comments on this manuscript. Dr. Kazushige Touhara (University of Tokyo) provided the mOR-EG-GFP mice, and Drs Peter Mombaerts (Max Planck Research Unit for Neurogenetics) and Tom Bozza (Northwestern University) provided the I7-IRES-tauGFP mice. We thank Dr. Michael Tordoff (Monell Chemical Senses Center) for proofreading of this manuscript’s draft.

